# Germline *APOBEC3B* deletion in Asian women increases somatic hypermutation in breast cancer that is associated with Her2 subtype, *PIK3CA* mutations, immune activation, and increased survival

**DOI:** 10.1101/2020.06.04.135251

**Authors:** Jia-Wern Pan, Muhammad Mamduh Ahmad Zabidi, Boon-Keat Chong, Mei-Yee Meng, Pei-Sze Ng, Siti Norhidayu Hasan, Bethan Sandey, Saira Bahnu, Pathmanathan Rajadurai, Cheng-Har Yip, Oscar M. Rueda, Carlos Caldas, Suet-Feung Chin, Soo-Hwang Teo

## Abstract

A 30-kb deletion that eliminates the coding region of *APOBEC3B* (*A3B*) is >5 times more common in women of Asian compared to European descent. This polymorphism creates a chimera with the *APOBEC3A* (*A3A*) coding region and *A3B* 3’UTR, and is associated with an increased risk for breast cancer in Asian women. Here, we explored the relationship between the *A3B* deletion polymorphism with tumour characteristics in Asian women. Using whole exome and whole transcriptome sequencing data of 527 breast tumours, we report that germline *A3B* deletion polymorphism leads to expression of the *A3A-B* hybrid isoform and increased APOBEC-associated somatic hypermutation. Hypermutated tumours, regardless of *A3B* germline status, were associated with the Her2 molecular subtype and *PIK3CA* mutations. Compared to non-hypermutated tumours, hypermutated tumours also had higher neoantigen burden, tumour heterogeneity and immune activation. Taken together, our results suggest that the germline *A3B* deletion polymorphism, via the *A3A-B* hybrid isoform, contributes to APOBEC-mutagenesis in a significant proportion of Asian breast cancers. In addition, APOBEC somatic hypermutation, regardless of *A3B* background, may be an important clinical biomarker for Asian breast cancers.

**Graphical Abstract:** 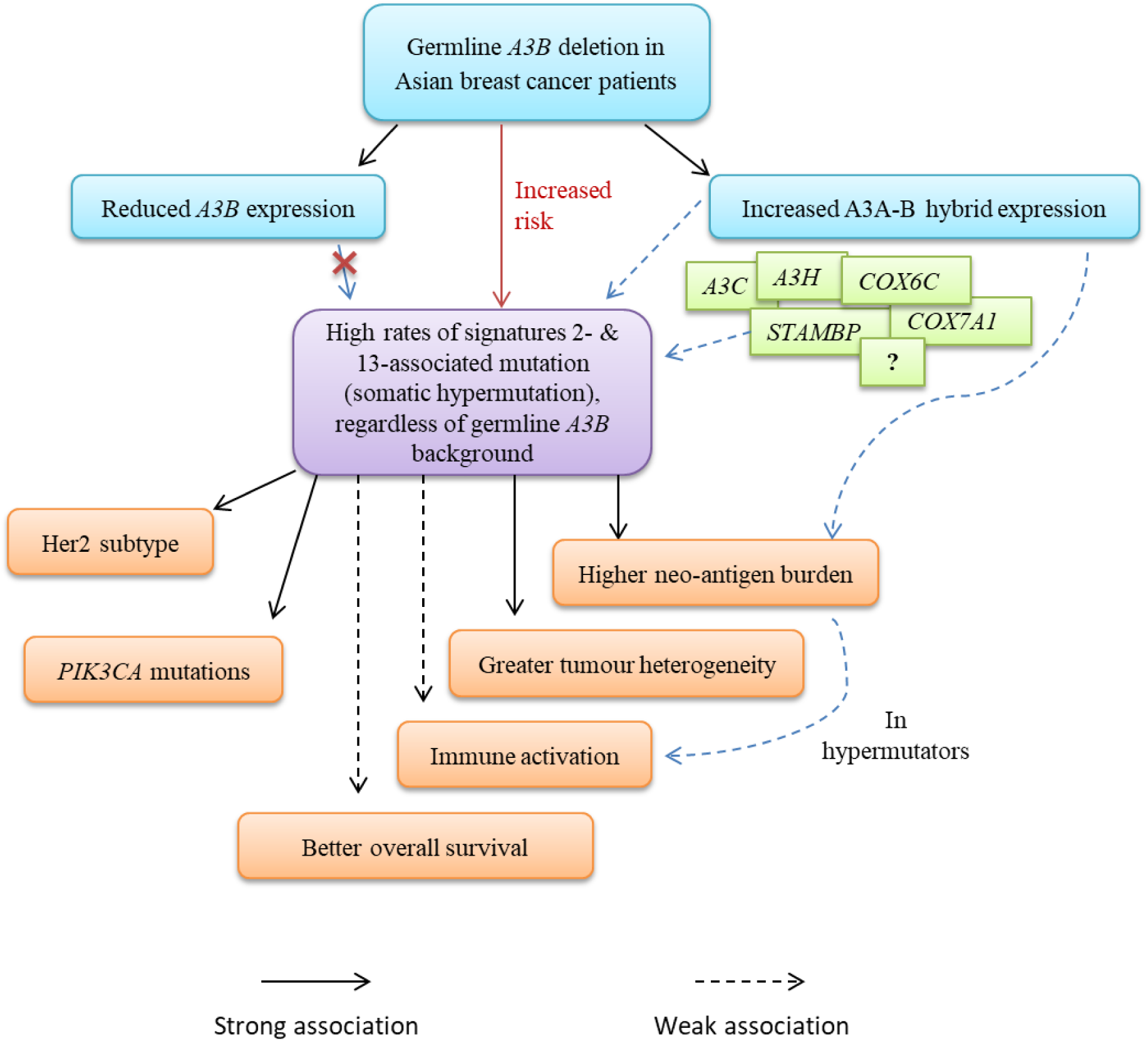

## Introduction

The identification of mutations in cancer cells driven by the APOBEC (“Apolipoprotein B mRNA- editing enzyme, catalytic polypeptide-like”) family of proteins^1^ has highlighted the role of intrinsic mutational processes in carcinogenesis and tumour evolution. APOBEC proteins are part of the AID/APOBEC superfamily of cytidine deaminases that include activation-induced deaminase (*AID*), *APOBEC1* (*A1*), *APOBEC2* (*A2*), *APOBEC3A-H* (*A3A* to *A3H*), and *APOBEC4* (*A4*), many of which play important roles in innate immunity^2,3^. More recently, *A3A* and *A3B*, and to a lesser extent, *A3H*, have been suggested to be endogenous sources of mutations for various cancers. Expression of *A3B* is associated with APOBEC-associated C-to-T transitions and is upregulated in breast, bladder, lung, head and neck, ovarian, and cervical cancer^4–7^. *In vitro* expression of *A3A* in cell lines has been shown to induce breaks in nuclear DNA and cell cycle arrest^8–10^. *A3H* has been postulated to contribute to mutagenesis in breast and lung cancer^11^, and *A3H* polymorphisms may contribute to risk for lung cancer^12^. Together, the data suggest that multiple APOBEC family members contribute to APOBEC mutagenesis in human cancers, with different effects in different cancers.

A common germline copy number polymorphism deleting a 30 kb coding region of the *A3B* coding sequence, fusing its 3’ UTR to the coding region of *A3A* resulting in an alternative *A3A-B* hybrid allele^13,14^ (Figure 1a), has been reported. In women of European descent, germline *A3B* deletion carriers in The Cancer Genome Atlas (TCGA) breast cancer cohort are twice as likely to develop cancers with a large number of mutations that correspond to APOBEC-driven mutational signatures^13^. The APOBEC mutational signature has been associated with neoantigen burden and immune activation^15–17^. This suggests that germline *A3B* deletion may generate tumour-specific antigens, which in turn activate the immune system in breast cancer patients. This may be mediated by the hybrid *A3A-B* gene isoform that is associated with APOBEC mutational signatures^18^, and may be more highly expressed in carriers of the germline *A3B* deletion^17^.

**Figure 1.**
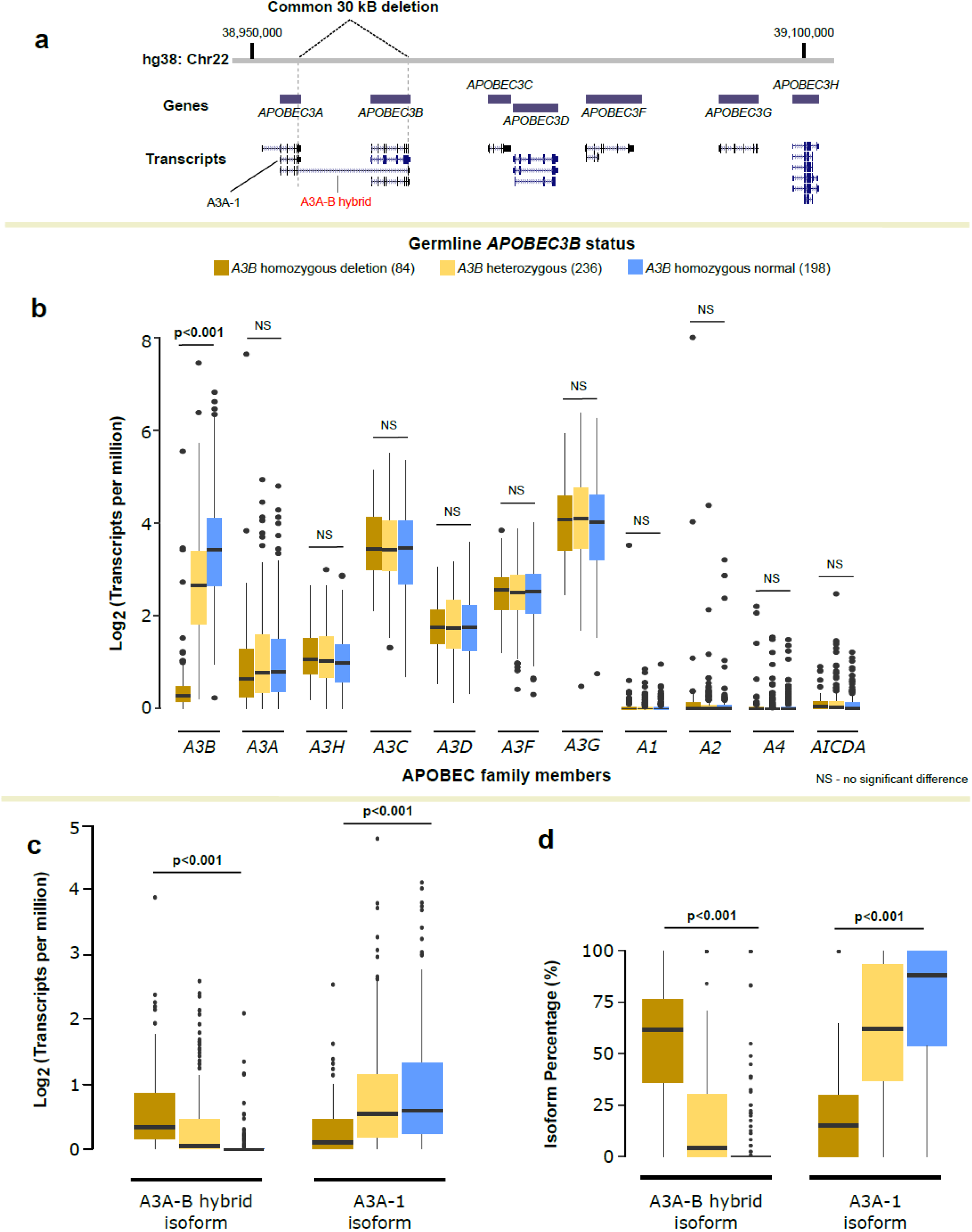
The relationship between germline *APOBEC3B* deletion and expression of APOBEC family members. **(a)** Diagram showing the APOBEC3 locus on chromosome 22, along with the transcript variants of each gene, modified from the UCSC Genome Browser. Location of a common germline deletion in the locus is also shown, where the deletion results in fusion of *A3A* coding sequence to the *A3B* 3’ UTR. **(b)** Gene expression of all APOBEC family members across *A3B* copy number, quantified as log transcripts-per-million (TPM). **(c)** Expression of the A3A-B hybrid isoform and the A3A-1 isoform in log TPM across *A3B* copy number. **(d)** Prevalence of the A3A-B hybrid isoform and A3A-1 isoform across different *A3B* copy numbers, measured as the percentage of total *A3A* gene expression. P-values indicated are for one-way ANOVA.

Intriguingly, there are population-specific differences in the APOBEC genomic locus. The deletion polymorphism is significantly more common in individuals of East Asian than of European ancestry (minor allelic frequency of 37% compared to 6%)^19^. This *A3B* deletion polymorphism is also associated with a modest increase in risk for breast cancer in women of Asian ancestry and in some studies of women of European ancestry, but not for European women in other studies^14,15,20–23^. These results raise the possibility that population-specific differences in genetic, lifestyle and environmental factors may modulate the impact of *APOBEC3B* deletion polymorphism in different populations. To investigate the biological consequences of germline *A3B* deletion on APOBEC mutagenesis and immune activation in the Asian population, we examined the effect of germline *A3B* deletion in a cohort of 527 breast tumours from Malaysia, where whole exome and whole transcriptomic sequencing information is available (Pan et al., in revision)^24^. The high prevalence of the polymorphism allowed for sufficient power to analyse homozygous and heterozygous carriers separately, and also enabled subtype-specific analyses.

## Results

### Germline *APOBEC3B* deletion results in reduced *APOBEC3B* expression and increased expression of an *APOBEC3A*-*B* hybrid isoform

With the transcriptomic data of 527 breast tumour samples from the MyBrCa cohort, we first investigated the relationship between germline *A3B* deletion and gene expression of the AID/APOBEC family members, particularly those located in the same genomic region (Figure 1a). As expected, homozygous *A3B* germline deletion carriers had 10-fold lower *A3B* expression than non-carriers (ANOVA, P < 0.001; Figure 1b), but there was no association between germline status and expression of other genes in the AID /APOBEC family. We also found that there was significantly higher *A3A-B* hybrid expression in germline *A3B* deletion carriers than in non-carriers, with a reciprocal loss of expression in the normal *A3A-1* transcript (Kruskal Wallis rank-sum test, P < 0.001; Figure 1c-d). In addition, we found that the overall expression of *A3A, A3B* and the A3A-1 and A3A-B hybrid isoforms were significantly different between different subtypes (highest in the basal subtype, lowest in the luminal A subtype) (Supp. Fig. 1), and germline *A3B* deletion leads to loss of expression of *A3B* and the *A3A-1* isoform, and an increase in expression of the *A3A-B* hybrid isoform, with the overall expression of *A3A* remaining similar in all breast cancer subtypes analysed (Supp. Fig. 1).

### Germline *APOBEC3B* deletion increases the risk for signature 2 and 13 hypermutation

Given that germline *A3B* deletion affects the expression of *A3B* and *A3A* isoforms and that these isoforms have been proposed to have different mutational activities^25^, we examined the relationship between germline *A3B* deletion status and mutational signatures of the breast tumours, focusing on the mutational signatures previously detected in breast tumours^26–28^. Of the mutational signatures analysed, tumours from germline *A3B*^del/del^ carriers were more likely to carry the APOBEC-associated Signatures 2 and 13 compared to non-carriers, while carriers of pathogenic variants in *BRCA1, BRCA2, PALB2, ATM* and *CHEK2* were more likely to have high levels of Signatures 3 (homologous recombination deficiency, HRD) than non-carriers (Fig. 2a-b). Although there was no significant difference in the number of total somatic mutations (single nucleotide variants (SNVs) + indels) across *A3B* deletion status (ANOVA, P = 0.347; Figure 2c), tumours in germline *A3B*del carriers have a higher proportion of mutations with Signatures 2 and 13, as well as a higher overall rate (mutational proportion multiplied by total somatic mutations) of Signature 2 and 13 mutations (Figure 2d-e). Notably, the level of mutational signatures 2 and 13 was highest in HER2-enriched breast tumours, but there was no association between germline *APOBEC3B* status and mutational burden in this subtype (Supp. Fig. 2). In contrast, we found that there was a significantly higher rate of mutational signatures 2 and 13 in germline *A3B* deletion carriers with luminal B breast cancers compared to non-carriers (Supp. Fig. 2).

**Figure. 2.**
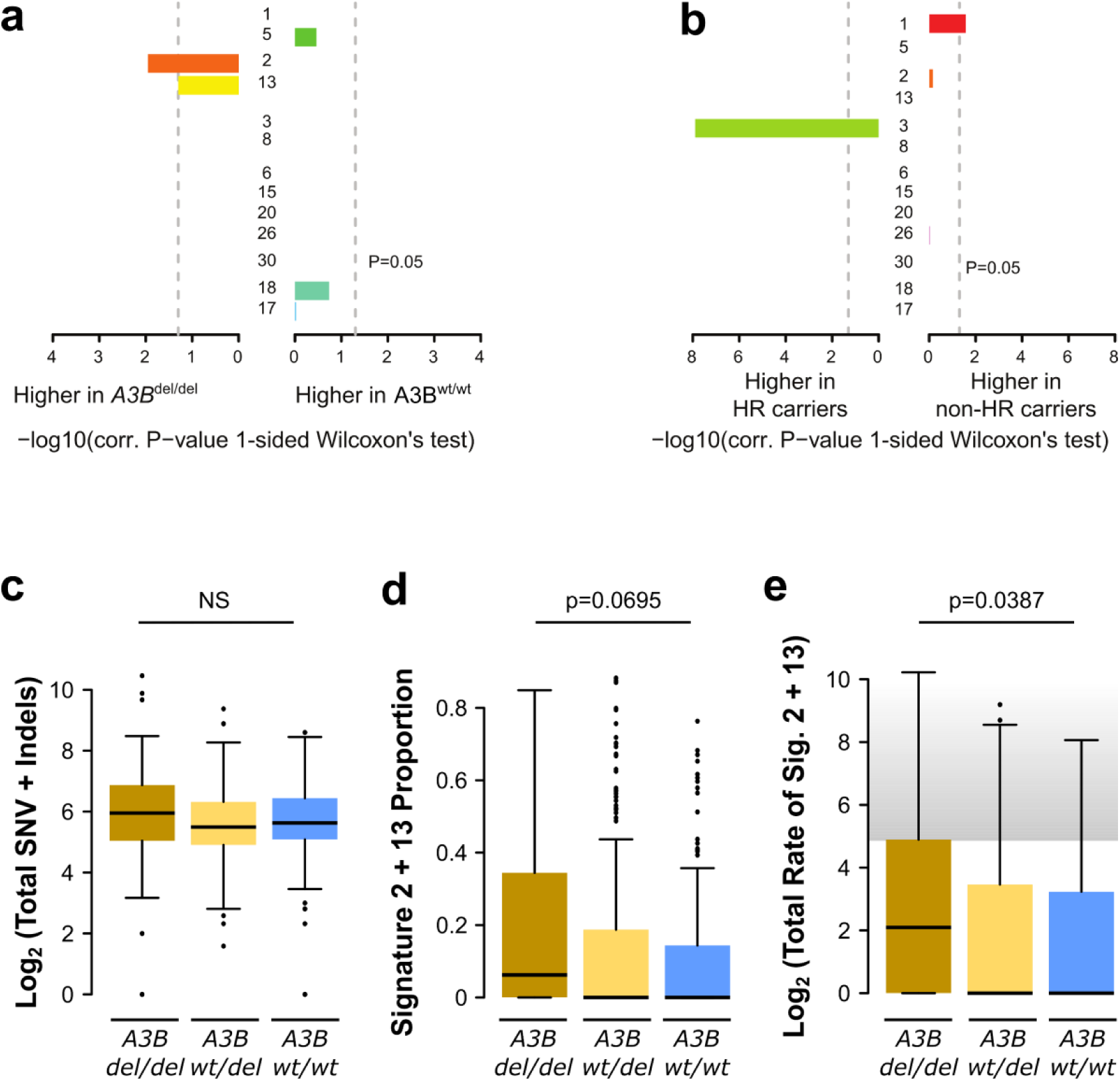
Germline *APOBEC3B* deletion and mutational signatures. **(a)** Comparison of mutational signatures between samples with homozygous germline *A3B* deletion and non-carriers, and, for contrast, **(b)** between samples with and without germline mutations in homologous recombination genes. **(c-e)** The (log-normalized) total mutational burden (c), the proportion of mutations with signatures 2 and 13 (d), and the (log-normalized) total rate of signature 2 and 13 mutations (e), between tumour samples with different germline *A3B* copy number. The grey area in (e) represents somatic hypermutation as defined by Nik-Zainal et al. (2014). P-values are for Kruskal Wallis rank sum tests (d) or one-way ANOVA (c,e).

We also found that Signature 2 and 13 (APOBEC) somatic hypermutation was significantly more common in germline *A3B* deletion carriers (Cochran-Armitage test for trend, P < 2.2 × 10^−16^; Supp. Table 1). Overall, individuals with one copy or two-copy deletion had a relative risk of 1.89 (95% confidence interval [CI]: [1.16, 3.06]) and 2.52 (95% confidence interval [CI]: [1.42, 4.45]) (Table 1) of bearing APOBEC hypermutated cancers. No association was observed with 3 other mutational signatures found in breast cancer (signatures 1, 3 and 8; Chi-squared test, P > 0.05, Supp. Table 2).

Next, we asked if signature 2 and 13 mutations were associated with *A3A* or *A3B* expression in breast cancer overall or in its subtypes. Overall, we found little association between total somatic mutation, the proportion of signature 2 and 13 mutations, and overall rates of signature 2 and 13 mutations with *A3B* gene expression (Supp. Fig. 3a). However, all three mutational measures were significantly associated with expression of both the A3A-1 and A3A-B hybrid isoforms (Supp. Fig. 3b-c). These associations appear to be strongest in the luminal B subtype, and weaker in the basal subtype (Supp. Fig. 4). Together, these results suggest that *A3A* may be a more important driver of signature 2 and 13 mutagenesis than *A3B*, and that the drivers of APOBEC mutagenesis may be slightly different across the various molecular subtypes.

To test the role of *A3A* in signature 2 and 13 mutagenesis, we compared the relative prevalence of mutations associated with *A3A* and *A3B* by examining the mutational context of all C→T transitions identified in our tumour samples through whole-exome sequencing (YT<u>C</u>A and RT<u>C</u>A for *A3A* and *A3B*, respectively^29^). We found that *A3A*-associated YT<u>C</u>A mutations were more common than *A3B*- associated RT<u>C</u>A mutations in the majority of our tumour samples, regardless of *A3B* deletion status or hypermutation (Supp. Fig. 5, Supp. Table 3). However, when the analysis was stratified by molecular subtype, this effect was observed in ER-positive and HER2-positive breast cancers, but appears to be weaker in the basal subtype (Supp. Fig. 6).

Finally, we conducted a multiple regression analysis to determine the factors associated with signature 2 and 13 mutational burden using a linear model. Using backward-stepwise elimination (see Methods), we obtained a minimal model in which only the *A3A-B* hybrid isoform, *A3C, A3H*, and three *A3A* interacting proteins (COX6C, COX7A1 and STAMPBP) were significantly associated with the total rate of signature 2 and 13 mutation (Supp. Fig. 7; Supp. Table 4). The minimal model also corresponded to the best model predicted using the Aikake information criterion (AIC), with an adjusted R-squared of 12.4%.

### Germline *APOBEC3B* deletion does not affect the molecular profile of breast cancers

To understand the impact of germline *A3B* deletion status on the molecular profile of breast cancer, we first examined at the frequency of the Integrative Cluster (IntClust) molecular subtypes across different *A3B* germline status. We found that there were no significant differences in the frequencies of molecular subtypes between *A3B* germline deletion carriers and non-carriers (Figure 3a), suggesting that germline *A3B* deletion does not significantly affect molecular subtype. We then examined the mutational profiles of these tumours by examining the non-synonymous mutations in the 10 most frequently affected genes in the MyBrCa cohort (Pan et al. 2020, submitted), comparing patients with different germline *A3B* copy number, but also did not find any significant differences (Figure 3b). We found that germline *A3B* deletion was marginally associated with a higher number of C-to-T or G-to-A transitions in *PIK3CA* (Supp. Fig. 8a), but not in the other nine genes. Comparison of *PIK3CA* hotspot mutations between samples with different *A3B* copy number revealed that the *PIK3CA*^E545K^ hotspot mutation is more common in carriers of the *A3B* deletion, while the *PIK3CA*^H1047R/L^ hotspot mutation is less common (Supp. Fig. 8b). Of note, the *PIK3CA*^E545K^ hotspot mutation is the result of the YT<u>C</u>A favoured mutational context for *A3A* mutagenesis.

**Figure 3.**
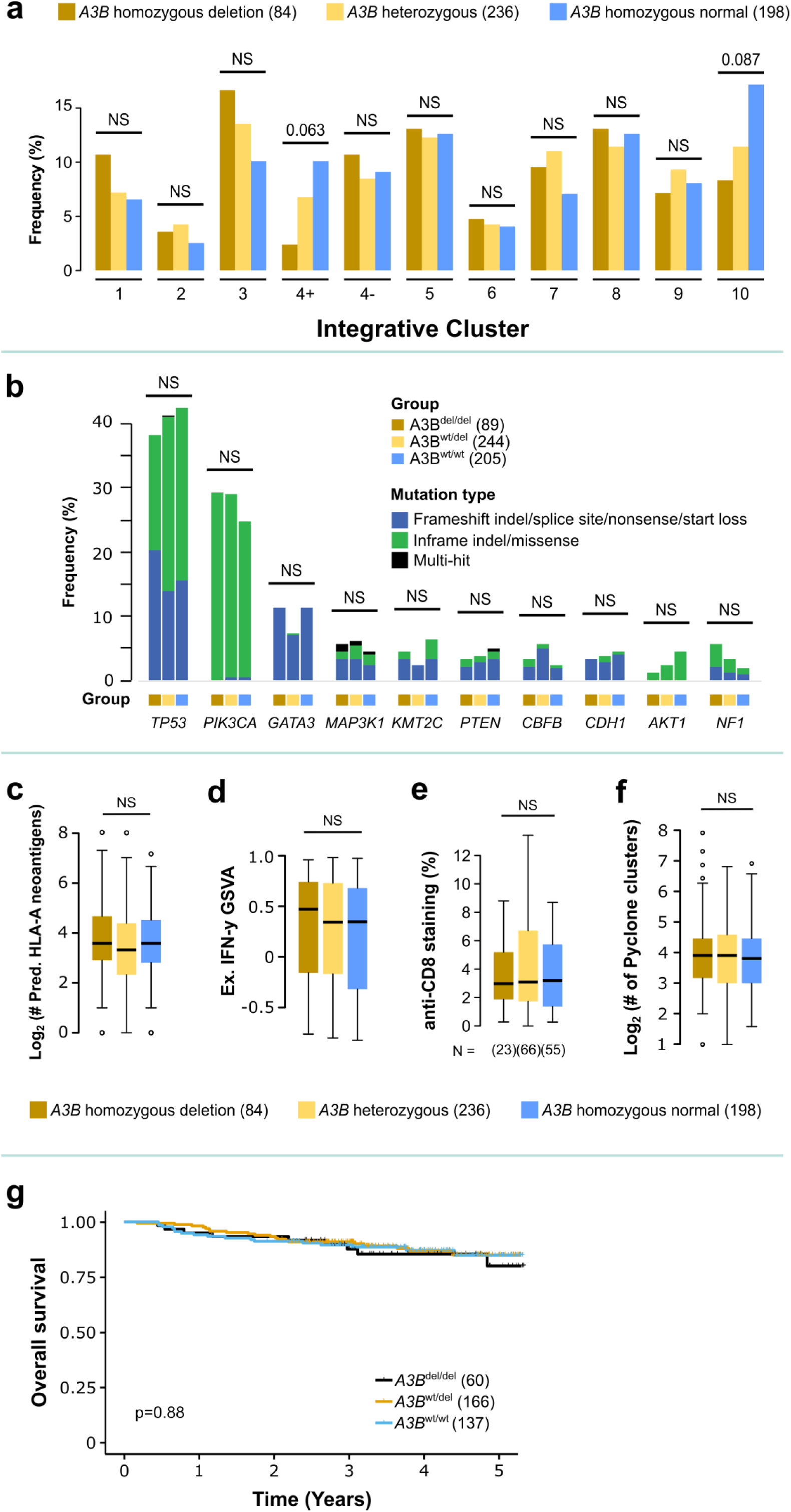
Molecular profiles of breast tumours with different germline *APOBEC3B* copy number. **(a)** Frequency of Integrative Cluster molecular subtypes across different *A3B* copy number. **(b)** Frequency of the top ten most commonly mutated driver genes across *A3B* copy number. (a-b) Sample sizes are indicated in brackets in the figure legend. Numbers above the bars are p-values for Fisher’s exact test. **(c)** Log-normalized counts of predicted HLA-A neoantigens in samples with different *A3B* copy number. **(d)** Gene set expression scores for the expanded IFN-γ gene set that is predictive of response to immunotherapy from Ayers et al. (2017), using the GSVA method, across *A3B* copy number. **(e)** Anti-CD8 IHC staining, measured as the percentage of area, of FFPE tumour samples with different *A3B* copy number. **(f)** Tumour heterogeneity, measured as the log-normalized counts of PyClone clusters, across different germline *A3B* copy number. (c-f) P-values indicated are for one-way ANOVA. **(g)** Kaplan-Meier plot of overall survival for patients with different *A3B* copy number. P-value indicated is for an unadjusted log-rank test.

As several recent reports have linked germline *A3B* deletion to the presentation of tumour-infiltrating immune cells^15,16^, we set out to explore if similar associations could be observed in our cohort. However, we found no significant difference in HLA-A neoantigen count across germline *A3B* deletion carriers (ANOVA, P = 0.1965; Figure 3c). Next, we characterised the immune scores of breast cancers across germline *A3B* copy number using four different algorithms, namely ESTIMATE^30^, GSVA^31^ using gene sets for immune cells from Bindea et al. (2013)^32^, GSVA using the expanded interferon-gamma (IFN-γ) gene set from Ayers et al. (2017)^33^ and the IMPRES method^34^. The IFN-γ and IMPRES methods have been used to predict response to checkpoint inhibitor immunotherapy in various cancer types. We did not find any significant difference across *A3B* copy numbers using the IFN-γ method (Figure 3d), or any of the other three algorithms (Supp. Fig. 9b). Immunohistochemistry (IHC) data using four different markers (anti-CD3, anti-CD4, anti-CD8, and anti-PD-L1) also revealed no significant difference in the staining for anti-CD8 (Figure 3e), as well as for anti-CD3, anti-CD4, and anti-PD-L1 (Supp. Fig. 10) across germline *A3B* deletion status. We further compared breast tumour heterogeneity, quantified using PyClone^35^, to *A3B* copy number, but observed no significant difference in tumour heterogeneity across germline *A3B* deletion (ANOVA, P = 0.3613; Figure 3f). Finally, our results also showed that *A3B* deletion status did not influence overall survival (Figure 3g), even though homozygous deletion carriers were significantly more likely to be Stage III, and significantly less likely to be Stage I (Supp. Table 5).

### APOBEC somatic hypermutation is associated with Her2 subtype, *PIK3CA* mutations, immune activation, and increased survival

Given our finding that germline *A3B* deletion increases the odds of developing APOBEC somatic hypermutation, we examined the relationship between APOBEC somatic hypermutation and the molecular profile of breast cancer in our cohort. To do so, we repeated the previous analyses, first by comparing the distribution of molecular subtypes and somatic mutations between samples with APOBEC hypermutation and samples without.

Interestingly, when we compared IntClust subtypes between APOBEC hypermutators and non-hypermutators, we found a higher prevalence of the Her2-associated IntClust 5 in the hypermutated tumours, with corresponding lower prevalence of IntClust 4- and IntClust 6 (Figure 4a). Tumours with APOBEC somatic hypermutation also had significantly higher (3-fold) *PIK3CA* mutations, marginally higher *KMT2C, CDH1*, and *NF1* mutations, and lower *GATA3* mutations (Figure 4b).

**Figure 4.**
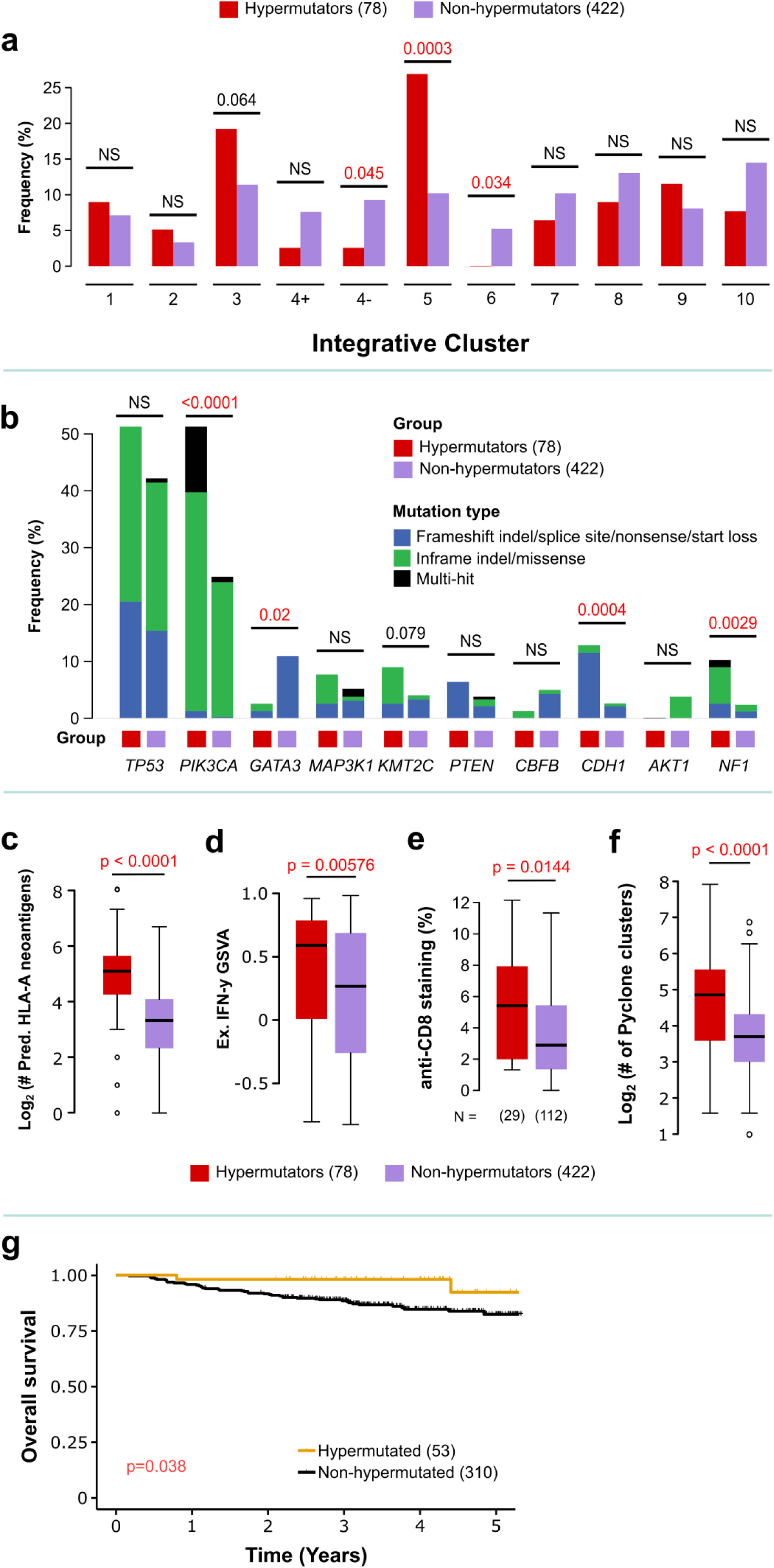
Molecular profiles of breast tumours with signature 2 and 13 (APOBEC) hypermutation. **(a)** Frequency of Integrative Cluster molecular subtypes in samples with and without APOBEC hypermutation. **(b)** Frequency of the top ten most commonly mutated driver genes in samples with and without APOBEC hypermutation. (a-b) Sample sizes are indicated in brackets in the figure legend. Numbers above the bars are p-values for Fisher’s exact test. **(c)** Log-normalized counts of predicted HLA-A neoantigens in samples with and without APOBEC hypermutation. **(d)** Gene set expression scores for the expanded IFN-γ gene set that is predictive of response to immunotherapy from Ayers et al. (2017), using the GSVA method, in samples with and without APOBEC hypermutation. **(e)** Anti-CD8 IHC staining, measured as the percentage of area, of FFPE tumour samples with and without APOBEC hypermutation. **(f)** Tumour heterogeneity, measured as the log-normalized counts of PyClone clusters, in samples with and without APOBEC hypermutation. (c-f) P- values indicated are for one-way ANOVA. **(g)** Kaplan-Meier plot of overall survival for patients with and without APOBEC hypermutation. P-value indicated is for an unadjusted log-rank test.

Next, when we compared neoantigen burden between APOBEC-hypermutated tumours and non-hypermutated tumours, we found that hypermutated tumours had a four-fold higher HLA-A neoantigen count compared to non-hypermutated tumours (ANOVA, P < 0.0001; Figure 4c). We found a weak association between HLA-A neoantigen count and *A3A-B* hybrid expression (Pearson’s correlation, *r* = 0.125, P = 0.0479; Supp. Fig. 9a), and a stronger association with the mutational rate of signatures 2 and 13 (Pearson’s correlation, *r* = 0.441, P < 0.0001; Supp. Fig. 9a). Furthermore, the association between APOBEC hypermutation and neoantigen burden was observed across all molecular subtypes (Supp. Figure 11).

Hypermutated tumours had slightly higher immune scores compared to non-hypermutators. This difference was most prominent when comparing IFN-γ immune scores (Figure 4d), although a similar trend also seen when comparing Bindea and ESTIMATE immune scores, but not with IMPRES scores (Supp. Fig. 9c). This finding was confirmed by IHC data, where hypermutators had a greater amount of staining than non-hypermutators for anti-CD8 (Figure 4e), as well as anti-CD3, anti-CD4, and anti- PD-L1 (Supp. Fig. 10). Additionally, comparisons of mutation rates to immune markers showed a modest correlation between signature 2 and 13 mutations and anti-CD3, anti-CD4, and anti-CD8 staining, supporting an association between APOBEC mutagenesis and the prevalence of tumour infiltrating lymphocytes in Asian breast cancer (Supp. Fig. 12). Notably, the association between immune scores and APOBEC hypermutation appears to be strongest in the luminal subtypes (luminal B in particular), with little to no association in ER- subtypes (Supp. Fig. 13)

We found that hypermutated tumours were predicted by PyClone to have two-fold more subclonal clusters than non-hypermutated tumours (Figure 4f). We also found a strong correlation between the rate of signature 2 and 13 mutation with tumour heterogeneity (Supp. Fig. 14), and the association between hypermutation and tumour heterogeneity was seen in all subtypes, albeit somewhat weaker in the basal subtype (Supp. Fig. 15).

Finally, we examined the association between APOBEC hypermutation and overall survival. Whilst there was no association between APOBEC hypermutated tumours and tumour stage (Supp. Table 5), patients with hypermutated tumours had better unadjusted overall survival than patients with non-hypermutated cancers (Figure 4g). Similarly, in a multivariable Cox proportional hazard model for overall survival that adjusted for germline A3B copy number, tumour stage, and subtype as covariables, APOBEC hypermutation had a hazard ratio of 0.4, albeit not statistically significant [0.1- 1.86] (Supp. Fig. 16). Overall, the data suggest that APOBEC hypermutation may be associated with better overall survival.

### Germline *A3B* deletion does not affect the molecular profile of tumours with APOBEC somatic hypermutation

Lastly, we asked if APOBEC somatic hypermutation associated with germline *A3B* deletion led to different outcomes than APOBEC somatic hypermutation driven by other sources. To test this, we focused only on tumours with APOBEC hypermutation and examined their molecular profiles in different *A3B* backgrounds. We found that the distribution of molecular subtypes and somatic mutations in APOBEC hypermutators were similar, regardless of *A3B* background (Figure 5a-b). Additionally, we found no significant difference in neoantigen burden (Figure 5c), immune scores (Figure 5d, Supp. Fig. 17), immune IHC markers (Figure 5e, Supp. Fig. 18), and tumour heterogeneity (Figure 5f, Supp. Fig 19) between APOBEC hypermutated tumours with different *A3B* backgrounds. Similar results were obtained in tumours without APOBEC hypermutation (Supp. Figs. 17-19). Overall survival of patients with hypermutation did not differ when they were further stratified by germline *A3B* deletion, and survival of patients with non-mutated cancers also did not differ by germline A3B status (Fig. 5g). Together, these results suggest that regardless of whether the source of the mutations is associated with germline *A3B* deletions, APOBEC hypermutation is associated with increased immune profiles and better survival.

**Figure 5.**
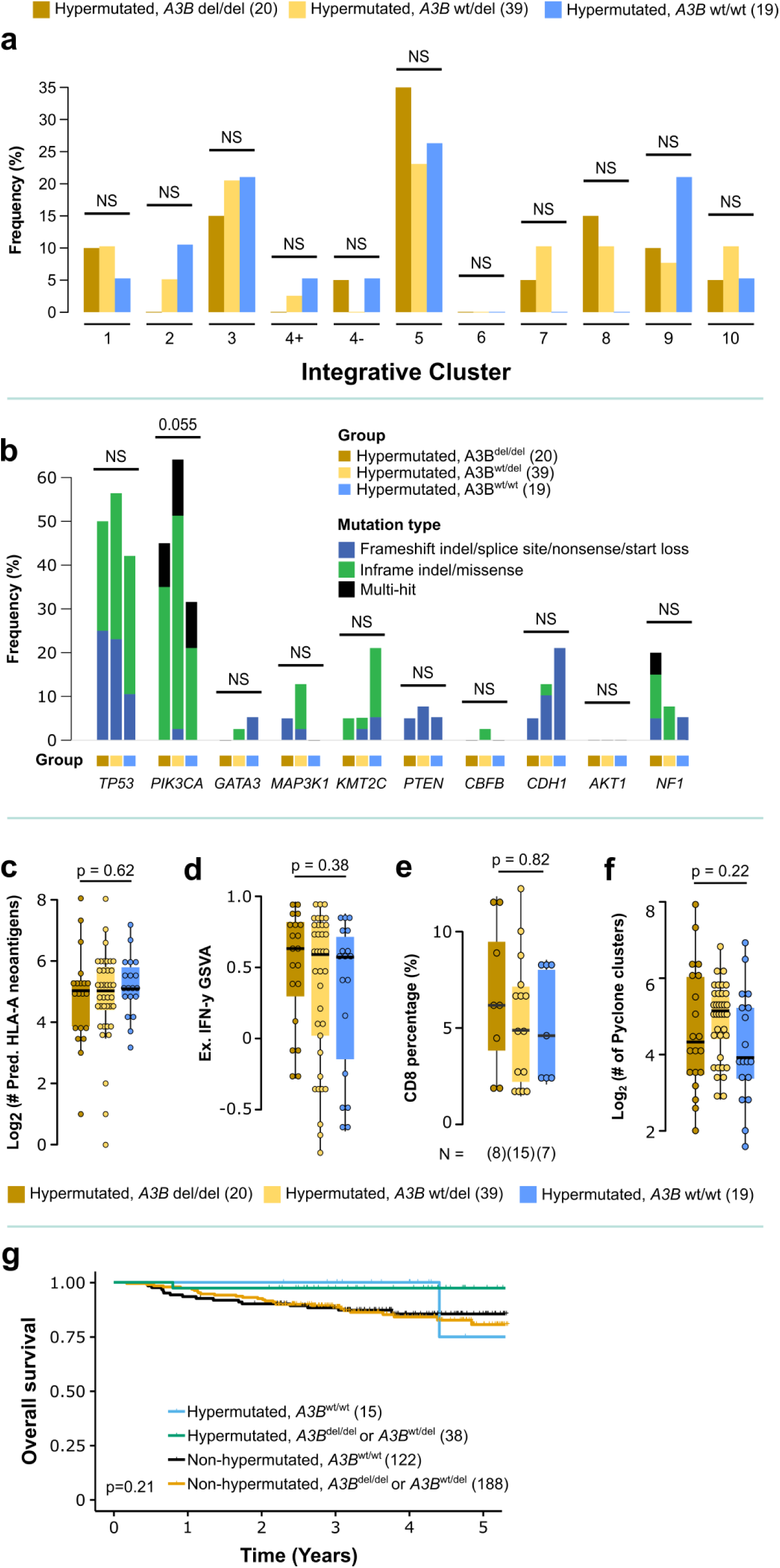
Comparison of molecular profiles of breast tumours with APOBEC hypermutation in different germline *APOBEC3B* copy number backgrounds. **(a)** Frequency of Integrative Cluster molecular subtypes in samples with APOBEC hypermutation across different *A3B* copy number backgrounds. **(b)** Frequency of the top ten most commonly mutated driver genes in samples with APOBEC hypermutation across different *A3B* copy number backgrounds. (a-b) Sample sizes are indicated in brackets in the figure legend. Numbers above the bars are p-values for Fisher’s exact test. **(c)** Log-normalized counts of predicted HLA-A neoantigens in samples with APOBEC hypermutation across different *A3B* copy number backgrounds. **(d)** Gene set expression scores for the expanded IFN-γ gene set that is predictive of response to immunotherapy from Ayers et al. (2017), using the GSVA method, in samples with APOBEC hypermutation across different *A3B* copy number backgrounds. **(e)** Anti-CD8 IHC staining, measured as the percentage of area, of FFPE tumour samples with APOBEC hypermutation, across different *A3B* copy number backgrounds. **(f)** Tumour heterogeneity, measured as the log-normalized counts of PyClone clusters, in samples with APOBEC hypermutation across different *A3B* copy number backgrounds. (c-f) P-values indicated are for one-way ANOVA. **(g)** Kaplan-Meier plot of overall survival for patients stratified by APOBEC hypermutation and *A3B* copy number. P-value indicated is for an unadjusted log-rank test.

## Discussion

In this study, we set out to investigate the biological consequences of germline *A3B* deletion on breast cancer in Asian women, where the deletion is more common compared to women of European descent. The high prevalence of the polymorphism allowed for sufficient power to analyse homozygous and heterozygous carriers separately, and to conduct subtype-specific analyses. We showed that germline *A3B* deletion results in reduced gene expression of *A3B* and the wild-type *A3A* isoform, with a reciprocal increase in expression of the *A3A-B* hybrid isoform. Heterozygous *A3B* deletion carriers have increased risk for APOBEC somatic hypermutation, particularly in the luminal B subtype. Importantly, we further showed that APOBEC somatic hypermutation, is associated with Her2-enriched molecular subtypes and *PIK3CA* mutations, higher levels of neoantigen burden, immune cell presentation, and tumour heterogeneity, and potentially better overall survival in Asian breast cancer patients. Furthermore, these associations seem to be independent of the source of mutation, as they were the same regardless of *A3B* background.

The association we found between APOBEC somatic hypermutation and immune infiltration may have potential implications for breast cancer immuno-oncology in populations where germline *A3B* deletion is common. Notably, our results are consistent with recent studies suggesting that APOBEC mutagenesis may identify individuals who benefit from immunotherapy^36–38^. On the other hand, our results are not entirely consistent with previous studies in tumour analysis of breast cancers in women of European descent that described a direct association between germline *A3B* deletion and immune activation^15–17^. One possibility is that the deletion only indirectly affects the immune microenvironment via APOBEC hypermutation, which is itself only weakly associated with expression of the *A3A-B* hybrid isoform, leading to a very weak overall association between germline *A3B* copy number and the immune microenvironment.

Our results raise an intriguing question of whether the association between APOBEC hypermutation and immune infiltration holds true for other cancer types in populations where the *A3B* deletion is common. An analysis of predominantly European TCGA pan-cancer data suggests the role of APOBEC mutagenesis in modulating the tumour immune microenvironment may be significant only in breast cancers^17^; on the other hand, whole-exome sequencing of 4000 Japanese tumours revealed a weak association between APOBEC mutational signatures and tumour mutational burden (itself a potential response biomarker for checkpoint inhibitors^39,40^) across several types of cancer^41^.

Our finding of a lack of association between germline *A3B* deletion and any specific molecular subtype is consistent with a previous report that found that the association between germline *A3B* deletion and increased risk for breast cancer is independent of ER status^20^. Likewise, our finding that APOBEC somatic hypermutation is highly associated with Her2-enriched molecular subtypes is evocative of previous reports that have linked Her2-enriched subtypes with higher *A3B* expression^42^. Besides that, the association between APOBEC mutagenesis with certain *PIK3CA* hotspot YT<u>C</u>A mutations has also previously been noted^43–45^, and may be related to the mutagenic predilections of the hybrid *A3A/B* isoform and *PIK3CA*’s role as an oncogene. Our results extend this association to germline *A3B* deletion as well, which may have important implications for therapies that target the PI3K pathway.

To our knowledge, this is the first study to suggest that APOBEC hypermutation modulates tumour heterogeneity and survival in Asian breast cancer. Some recent reports have indicated that *APOBEC* mutagenesis contributes to tumour heterogeneity in late stage non-small cell lung carcinoma (NSCLC) and metastatic thoracic tumours^46,47^, but its role in breast cancer was previously not analysed. The link between APOBEC hypermutation and tumour heterogeneity is unsurprising given that mutagenesis should tend to increase cellular diversity. Likewise, the positive survival effect we found is consistent with previous studies on Asian oral cancer^48^, and may be the result of hypermutators having higher numbers of tumour-infiltrating immune cells^38^, which has been linked to better overall survival in some subtypes of breast cancer^49^. On the other hand, our results contradict previous studies that suggest that APOBEC mutagenesis is associated with poor prognosis in other cancers, such as multiple myeloma^50^. Our study also confirms previous reports which found that, in contrast with APOBEC mutagenesis or *A3A*/*A3B* mRNA expression, germline *A3B* copy number does not have prognostic value^23,51^.

To our knowledge, this is also the first study to elucidate the subtype-specific associations of germline *A3B* deletion and APOBEC mutagenesis in breast cancer. Our finding that germline *A3B* deletion is a stronger driver of APOBEC mutagenesis in the luminal B subtype, but a weaker driver in the basal subtype, suggests that further research in this area will need to account for breast cancer subtype as a significant variable. Similarly, our finding that APOBEC hypermutation is more strongly associated with immune activation in the luminal B subtype may help to focus future research efforts.

Multiple lines of evidence from our analysis suggest that in *A3B* deletion carriers, increased expression of the *A3A-B* hybrid isoform drives mutagenesis, suggesting that the *A3B* 3’-UTR may play an important role in modulating APOBEC mutagenesis. Together with previous studies^25,29^, this finding contributes to the resolution of the paradox that loss of *A3B*, an endogenous mutator, is associated with an increased risk of breast cancer in the Asian population by supporting the hypothesis that the *A3A-B* hybrid isoform is a more potent mutator than either the *A3A* or *A3B* normal isoforms. It is also interesting to note that the three *A3* members that are significantly associated with APOBEC mutagenesis in our linear model each contain only one zinc-dependent cytidine deaminase domain (ZD-CDA) and exhibit both cytoplasmic and nuclear localisations^52^. While it is conceivable that *A3D, A3F* and *A3G* did not appear in the model (they are cytoplasmically localised and are even excluded from chromatin throughout mitosis hence have no contact with nuclear DNA^53^), it is not immediately clear why *A3B*, which has access to nuclear DNA and is widely regarded as the endogenous source of mutation^5–7,42,54^, did not turn out as a significant predictor in our model. This may or may not be related to the recent finding that APOBEC3 mutagenesis *in vitro* is episodic, occurring in intermittent and irregular bursts^55^.

As a whole, our study in Asian women validates the results from previous analyses on the biological consequences of germline *A3B* deletion on breast cancer that were conducted predominantly in women of European descent. In particular, our study highlights the significance of APOBEC hypermutation as an important predictor of the breast tumour microenvironment, and suggests that Asian breast cancer patients with APOBEC hypermutation may be more amenable to immunotherapy.

## Acknowledgements

This project was funded by a research grant from the Newton-Ungku Omar Fund (MRC Ref: MR/P012442/1) by the British Council and the Malaysian Industry-Government Group for High Technology (MIGHT) to CC and SHT. Cancer Research Malaysia also receives charitable funding from the Scientex Foundation, Estée Lauder Companies, Yayasan Petronas, and Yayasan Sime Darby which contributed to the funding of this study. OMR, CC, and SFC also receive funding from Cancer Research UK. The authors would like to thank Dr. Tan Min Min and Nadia Rajaram for help with data curation, nurses and staff who helped with sample collection, as well as Tan Wee Lin, and all staff at the Subang Jaya Medical Centre Tissue Diagnostics laboratory for assistance with histopathological sample retrieval and processing. All genomics work was undertaken by the Genomics Core Facility CRUK Cambridge Institute.

## Author Contributions

JWP led the data analysis and wrote the manuscript. MMAZ and BKC contributed to data analysis and helped to draft the manuscript. MYM, PSN, SB, SNH, and CHY contributed to sample collection and processing and data collection, while BS, OMR, and SFC generated and collected data. PR provided histopathology expertise, and together with CHY collected clinical data. OMR, CC, SFC, and ST designed experiments, interpreted results, and drafted the manuscript. The project was directed and co-supervised by OMR, CC, SFC and ST, and were responsible for final editing.

## Methods

### Data Description

Data for this project was taken primarily from the MyBrCa cohort tumour sequencing project. In brief, shallow whole-genome sequencing (sWGS), whole-exome sequencing (WES), and RNA-sequencing were conducted on 560 sequential female breast cancer patients from a single hospital in Subang Jaya, Malaysia, and analysed together with available clinical and overall survival data. The cohort data and sequencing methods are described in full in Pan et al. (in revision)^24^. Only the methods important or unique to this study are described below.

### Determining the Presence of the *APOBEC3B* Deletion

To determine the presence of the *APOBEC3B* (*A3B*) germline deletion for each individual, we calculated for each matched normal WES sample the ratio (*r*) of the mean depth of coverage for 35 loci in exons within the deletion (*d*_in_) to the mean depth of coverage for 70 loci in exons immediately adjacent to the deletion (*d*_out_). The list of loci used is identical to the list used in Nik Zainal et al. 2014^13^. Then, we designated each sample as being homozygous, heterozygous, or normal for the *A3B* deletion (*A3B*^del/del^, *A3B*^wt/del^, and *A3B*^wt/wt^ respectively) by fitting a 3-component mixture model to the distribution of *r* using an expectation-maximization algorithm as implemented in the mixtools (v. 1.1) package in R. The gene diagram of the APOBEC locus in Figure 1 is adapted from the UCSC Genome Browser.

### Molecular Subtyping

We transformed gene-level count matrices for the MyBrCa cohort into log_2_ counts-per-million (logCPM) using the voom function from the limma (v. 3.34.9) R package. We then performed quantile normalization on each individual transformed matrix, followed by subtyping according to PAM50 and SCMgene designations using the Genefu package in R (v. 2.14.0), and according to integrative clusters using the iC10 R package (v. 1.5).

### Mutational profiling

For SNVs, we used positions called by Mutect2 with following filters: minimum 10 reads in tumour and 5 reads in normal samples, OxoG metric less than 0.8, variant allele frequency (VAF) 0.075 or more, p-value for Fisher’s exact test on the strandedness of the reads 0.05 or more, and S_AF_ more than 0.75. For positions that are present in 5 samples or more, we removed two positions that were not in COSMIC and in single tandem repeats. We also removed variants that have VAF at least 0.01 in gnomAD, and considered only variants that are supported by at least 4 alternate reads, with at least 2 reads per strand. For indels, we also required the positions to be called by Strelka2. Variants were annotated using Oncotator version 1.9.9.0. PIK3CA lollipop plots were generated using the MutationMapper tool from cBioPortal^56^.

### Mutational Signatures

To determine the prevalence of previously-reported breast cancer mutational signatures from COSMIC matrices (Signatures 1, 2, 13, 3, 8, 6, 15, 20, 26, 5, 17, 18 and 30), we used deconstructSigs^57^, restricted to samples with at least 15 SNVs. To determine the difference in mutational signature weights between *A3B*^del/del^ and *A3B*^wt/wt^ germline carrier samples, we performed 2-sided rank sum Wilcoxon’s test on the two categories. Rates of mutational signatures 1, 2, 3, 8 and 13 were calculated by multiplying the total somatic SNVs and indels with the proportions of each mutational signature.

### Identification of Hypermutators

Using the combined rates of mutational signatures 2 and 13 (simple addition of the rates of each signature), hypermutators were identified based on Nik Zainal et al.’s definition of hypermutators as samples that had a mutational rate of signatures 2+13 exceeding 1.5 times the interquartile range from the 75^th^ percentile^13^. Outliers for other mutational signatures were identified using a similar approach.

### Profiling the Tumor-Immune Microenvironment

We assessed overall immune cell infiltration in the bulk tumour samples from RNA-seq TPM gene expression scores using ESTIMATE (v. 1.0.13)^30^, as well as with GSVA (v. 1.26) using the combined immune cell gene sets from Bindea et al. 2013^32^. We also scored each sample for the immune features predictive of checkpoint inhibitor immunotherapy using IMPRES scores^58^, as well as GSVA using the Expanded IFN-gamma gene set from Ayers et al. 2017^33^.

### Validation of Immune Scores with Immunohistochemistry

To quantify tumour infiltrating lymphocytes, FFPE blocks for 154 patients with sequencing data were sectioned and stained for anti-CD3 (clone 2GV6, predilute; Ventana Medical Systems), anti-CD4 (clone SD35, predilute; Ventana Medical Systems), anti-CD8 (clone SD57, predilute; Ventana Medical Systems) and anti-PD-L1 (clone SP263, predilute; Ventana Medical Systems) using an automated immunostainer (Ventana BenchMark ULTRA; Ventana Medical Systems, Tucson, AZ). Stained slides were digitized using an Aperio AT2 whole slide scanner. CD3, CD4, and CD8 staining was quantified using the Aperio Positive Pixel digital pathology tool (v9 algorithm at 0.16 colour saturation). PD-L1 expression was determined using the Combined Positive Score system.

### Quantification of Tumour Heterogeneity

Tumour heterogeneity was determined using PyClone (v 0.13.1)^35^ with default options to estimate the number of subclonal clusters within each tumour sample. Allele counts used for the PyClone input were extracted from the GATK output MAF files, while copy number input data was generated by ASCAT (v. 2.5.2) from WES allele counts generated by alleleCounter (v 4.0.1).

### Survival analysis

For each patient, overall survival data was obtained by querying their names and identity card numbers against the Malaysian National Registry records of deaths. Patients that did not return any matches against the database were assumed to still be alive, and *vice versa*. Length of survival was defined as the period of time from the date when patients were recruited into the study until the date of death as recorded by the Malaysian National Registry for patients who have passed away, or until the date when the Malaysian National Registry was last queried for patients assumed to still be alive. For all survival analyses in this study, only patients with at least two years of survival data were included (n = 367). Unadjusted Kaplan-Meier analyses and log rank tests were conducted using the survival package in R (v. 2.44) and plotted using the “ggtsurvplot” function from the survminer R package (v. 0.4.4). Cox proportional hazard models were built using the “coxph” function from the survival package and plotted using the “ggforest” function from survminer.

### Neoantigen Analysis

Sample HLAs were determined using Polysolver^59^ from tumor and normal DNA WES data. Only HLA alleles that were concordant in tumor and normal WES data were considered. In this analysis, we focused only on HLA-A. This selection was made because the majority of studies identifying peptides that bind to HLA molecules have focused on those recognised by cytotoxic T lymphocytes, hence the prediction of antigen binding to MHC class I molecules is the most studied^60^. Amongst MHC class I, HLA-A is ranked among the genes with the fastest evolving coding sequence as a result of ever-changing antigen selection pressures, thus serving as a good reflection of neo-antigen burden. Somatic mutations were annotated using the Ensembl Variant Effect Predictor. All possible neoantigen peptides (9- to 11-mers) encompassing the nonsynonymous mutations were predicted using a combination of NetMHCpan and NetMHC on the pVAC-seq platform. Only neoantigens with predicted binding of less than 500nM were considered.

### Extraction of Isoform Data

Raw RNA-Seq reads were mapped to the GrCh38 reference human genome, and isoform-level expression was quantified as transcripts-per-million (TPM) using RSEM (v. 1.2.31) with the *Homo sapiens* GRCh38.97 genome annotation model.

### Gene Expression Statistical Analyses

The difference in transcript per million (TPM) expression of all APOBEC family members across *A3B*^del/del^, *A3B*^wt/del^, and *A3B*^wt/wt^ breast cancers was compared using analysis of variance (ANOVA) whereas difference in isoform TPM and percentages were compared using the Kruskal Wallis rank sum test (Figure 1) as these data were not normally distributed. As for the total somatic mutation (SNVs and indels) and rates of mutational signatures 2 and 13 across A3B deletion status, the difference was tested using ANOVA while Kruskal Wallis test was used for the total proportion of mutational signatures (Figure 3). The association between somatic mutations, proportions and rate of signatures 2 and 13 with A3B and A3A-B hybrid expression was tested using Spearman’s regression. As for HLA-A neo-antigen count and tumour heterogeneity, logarithmic transformation of these data was normally distributed, hence Pearson’s correlation was applied in these cases. The difference across A3B deletion and hypermutation status for these data was compared using ANOVA whereas those for immune scores was tested using Kruskal Wallis and Wilcoxon rank sum tests, respectively, for three- and two-group comparisons.

### Backward-stepwise elimination for linear modelling and Akaike Information Criterion

The minimal model was built from the original global model using backward-stepwise elimination method implemented in the MuMIn package in R. In our first constructed linear model, the predictors consisted of all APOBEC family members. Subsequent models involved addition of known A3A interacting proteins as predictors and the interaction terms between A3A and its interacting proteins, hoping to improve the explanatory power of our model. The minimal models were then compared to all the possible alternative models generated by AIC to generate a final model.

## Supplemental Figures

**Supp. Fig. 1.**
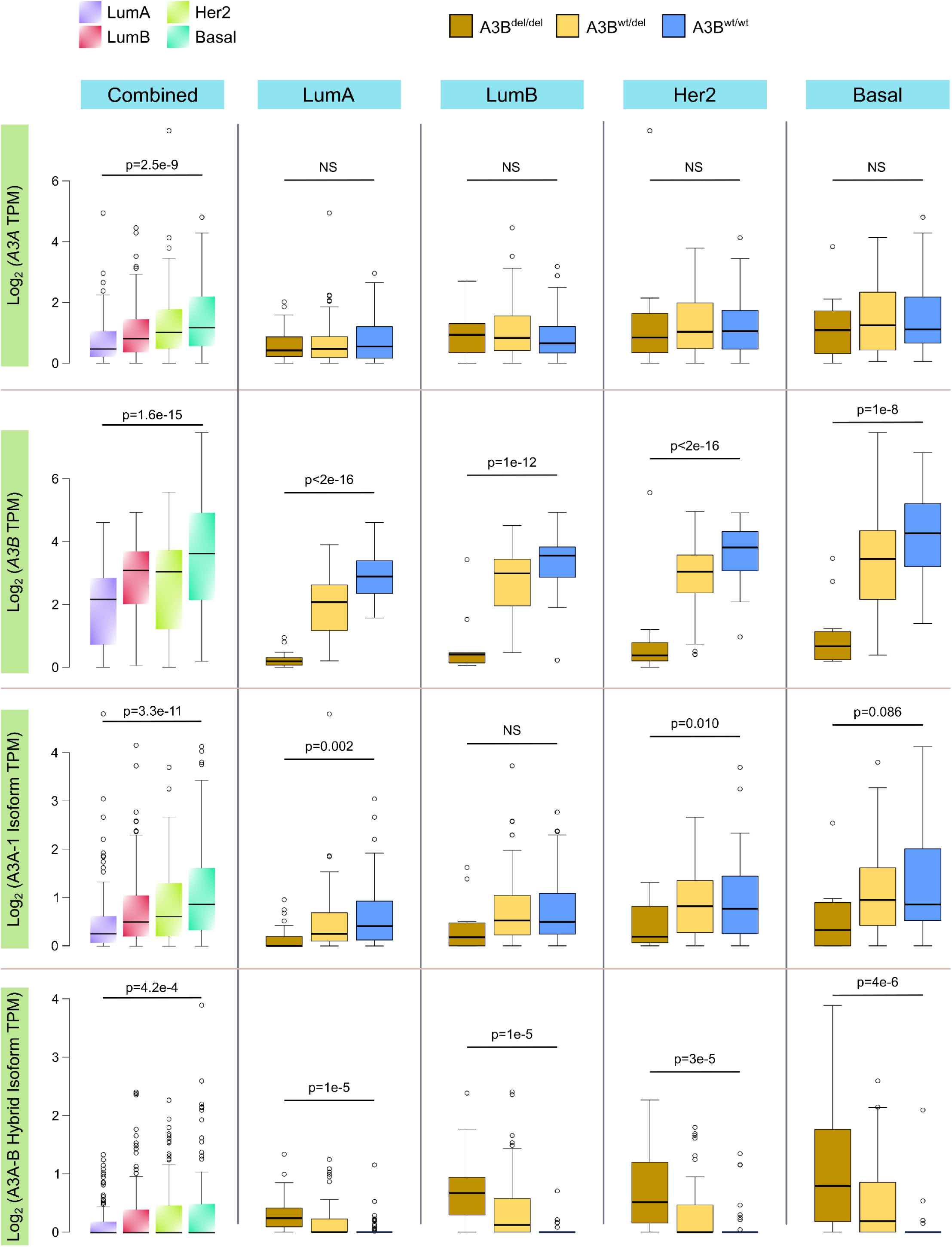
APOBEC expression of across breast cancer subtypes. Expression of the *A0POBEC3A* and *APOBEC3B* genes, as well as the A3A-1 and A3A-B hybrid isoforms of *A3A*, across the different PAM50 breast cancer molecular subtypes as well as germline *A3B* copy number, quantified as log transcripts-per-million (TPM). P-values indicated are for one-way ANOVA.

**Supp. Fig. 2.**
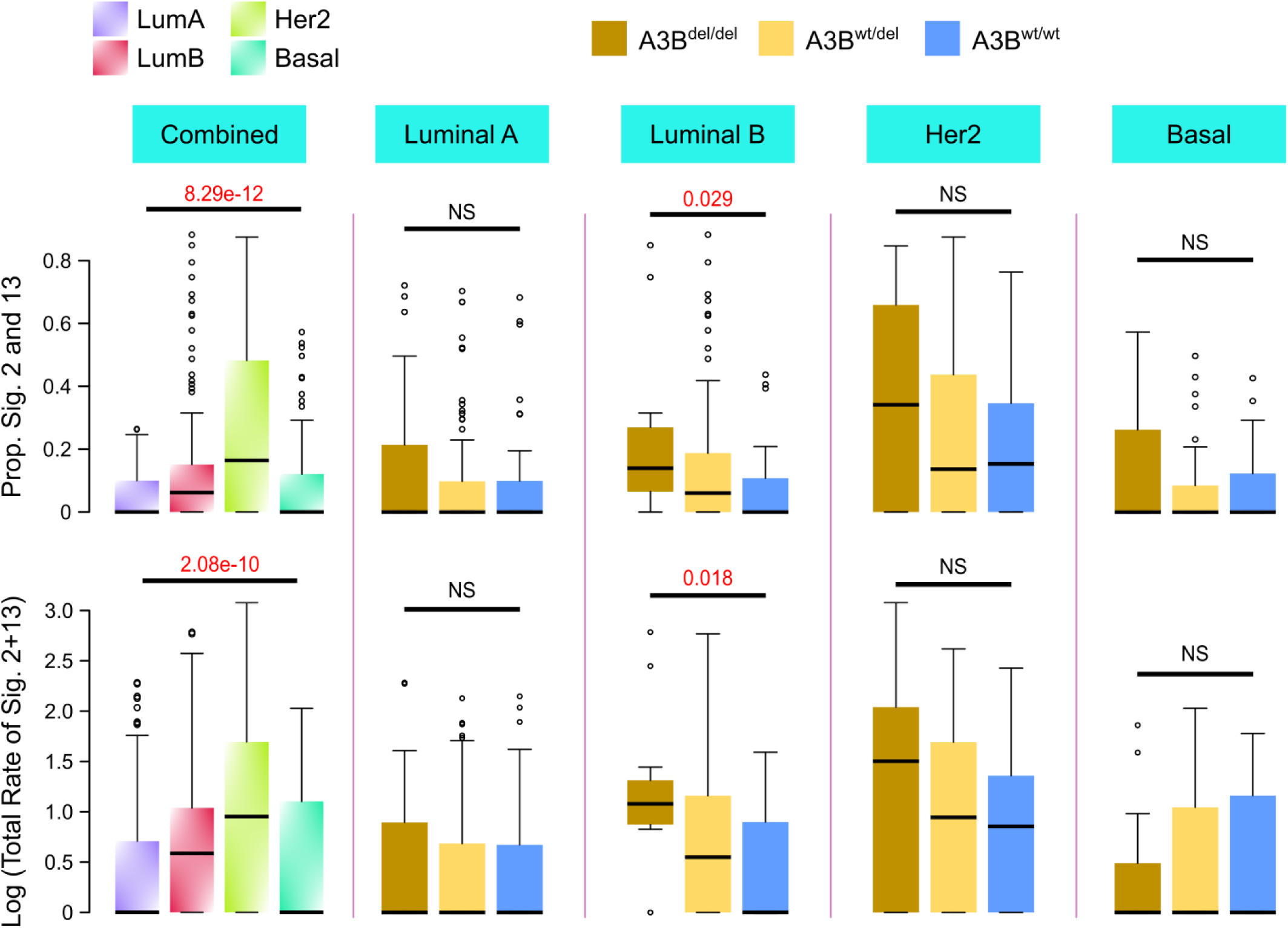
Germline *APOBEC3B* deletion and mutational signatures 2 and 13 across subtypes. Comparison of the proportion and total rate of mutational signatures 2 and 13 between samples with different germline *A3B* copy number, stratified by PAM50 molecular subtype. P-values shown are for one-way ANOVA.

**Supp. Fig. 3.**
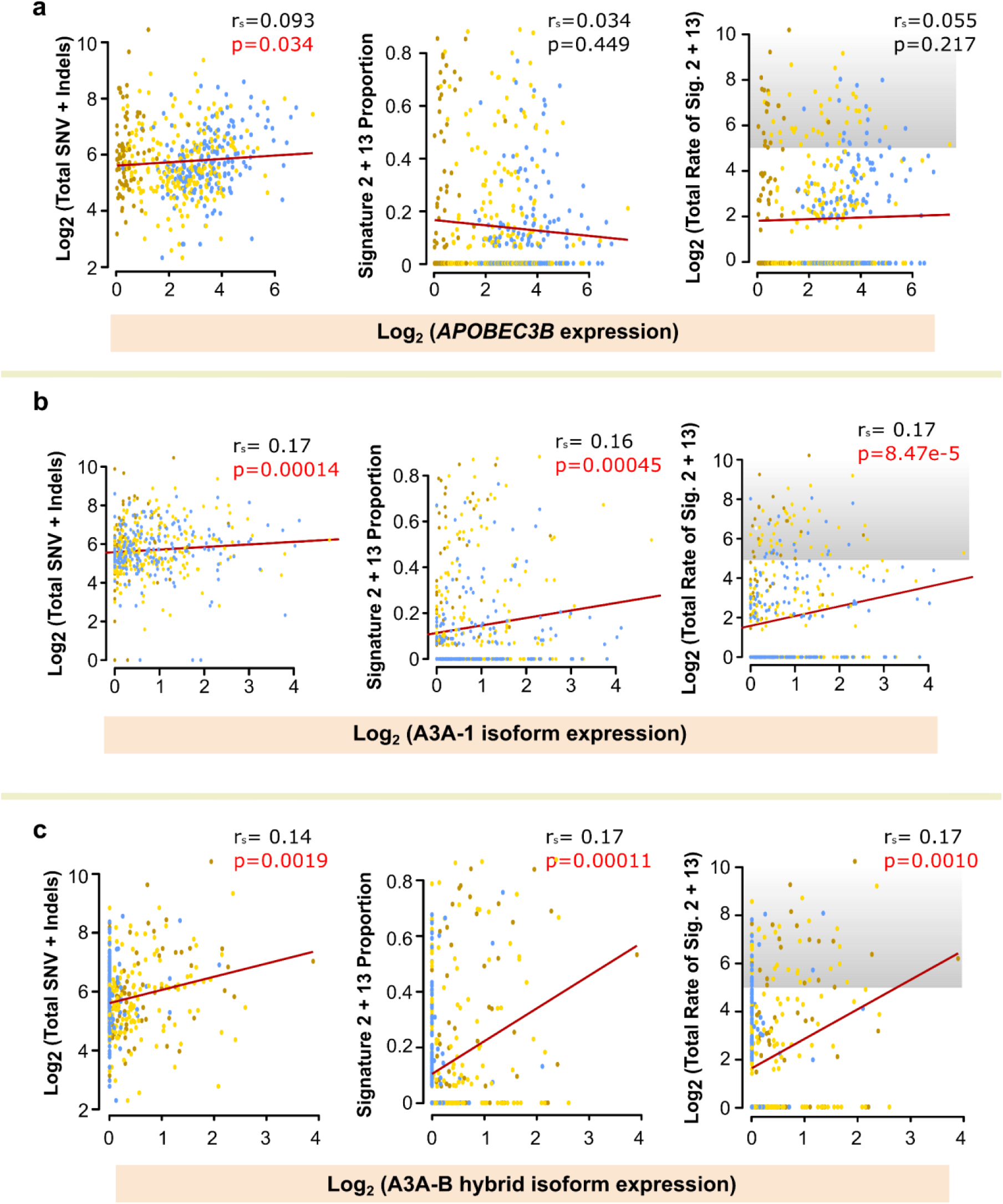
Relationship between APOBEC expression and signature 2/13 mutations. **(a)** Comparison of total mutational burden (left), the proportion of mutations with signatures 2 and 13 (middle), and the total rate of signature 2 and 13 mutations (right) to expression of the *APOBEC3B* gene. **(b-c)** Comparison of total mutational burden (left), the proportion of mutations with signatures 2 and 13 (middle), and the total rate of signature 2 and 13 mutations (right) to expression of the (b) A3A-1 and (c) A3A-B hybrid gene isoforms. In the right-sided figures, the grey area represents APOBEC somatic hypermutation as defined by Nik Zainal et al. 2014.

**Supp. Fig. 4.**
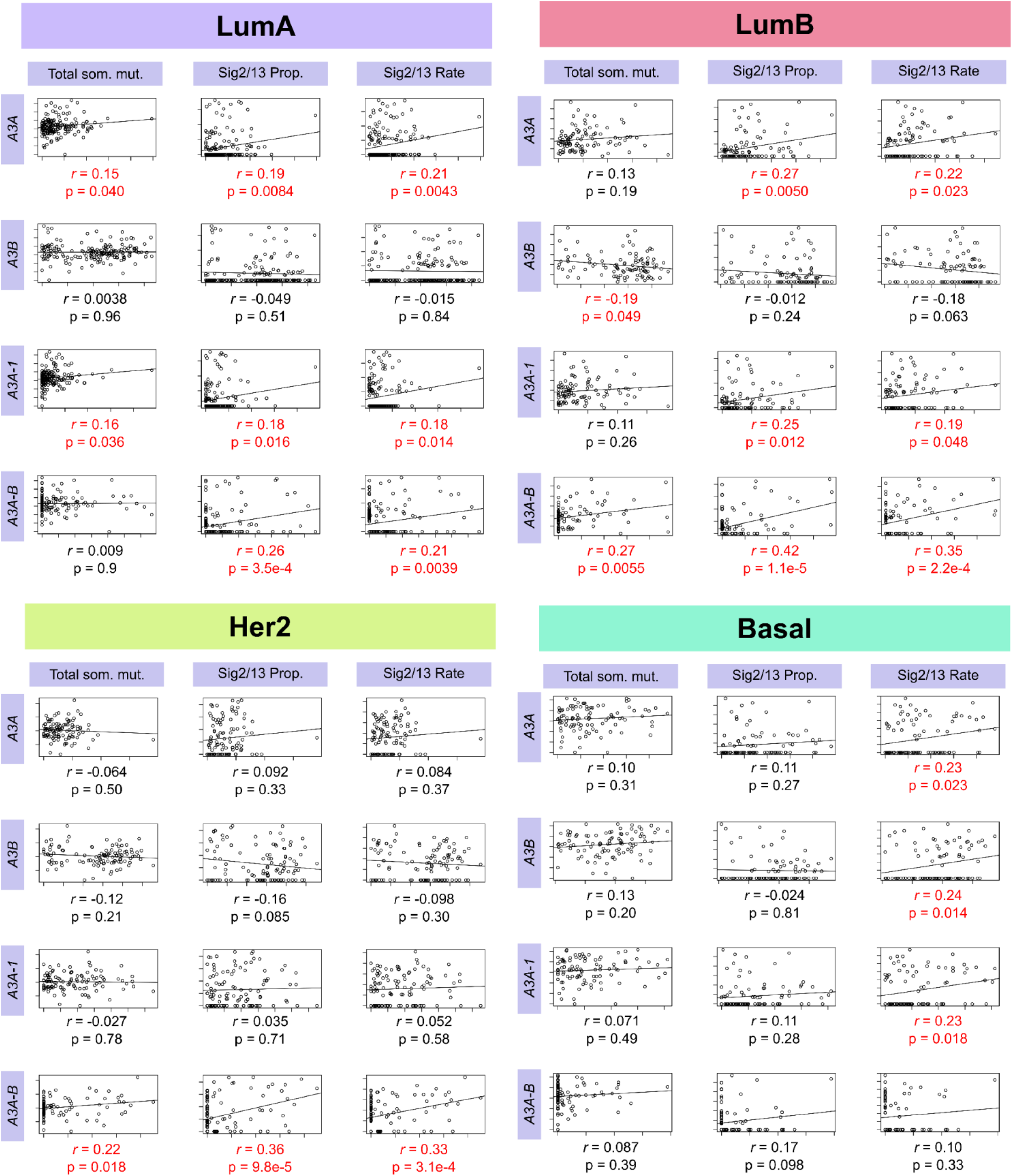
Relationship between APOBEC expression and signature 2/13 mutations in different molecular subtypes. Comparison of log-normalized total mutational burden (left), the proportion of mutations with signatures 2 and 13 (middle), and the log-normalized total rate (proportion of mutations multiplied by total mutational burden) of signature 2 and 13 mutations (right) to expression of the *APOBEC3A* (top) and *APOBEC3B* (second from top) genes, as well as expression of the A3A-1 (second from bottom) and A3A-B hybrid isoforms, for each main PAM50 molecular subtype. Gene/isoform expression is quantified as log2-normalized transcripts-per-million. Also indicated are the Pearson’s correlation coefficient (*r*) for each comparison, and the corresponding p-value. Comparisons with a p-value below 0.05 are highlighted in red.

**Supp. Fig. 5.**
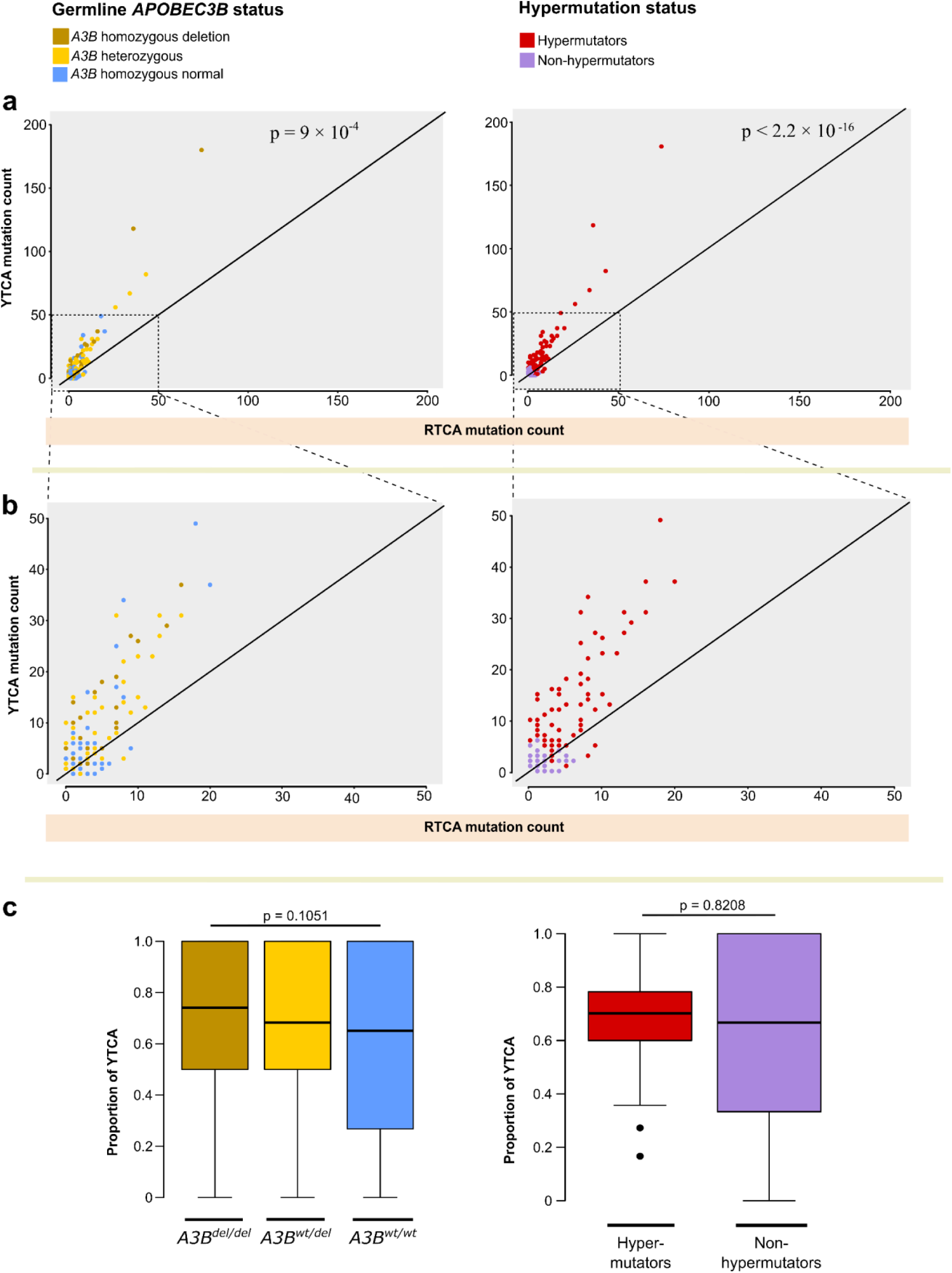
Comparison of RT<u>C</u>A and YT<u>C</u>A mutations. **(a)** Comparison of the number of YTCA mutations to the number of RTCA mutations in each MyBrCa sample. Each sample is colored according to germline *A3B* status (left) or hypermutation status (right). Black line indicates a 1:1 ratio. **(b)** Zoomed-in version of (a), for samples with mutation counts of less than 50. **(c)** Prevalence of YTCA mutations across *A3B* copy number (left) and hypermutation status (right).

**Supp. Fig. 6.**
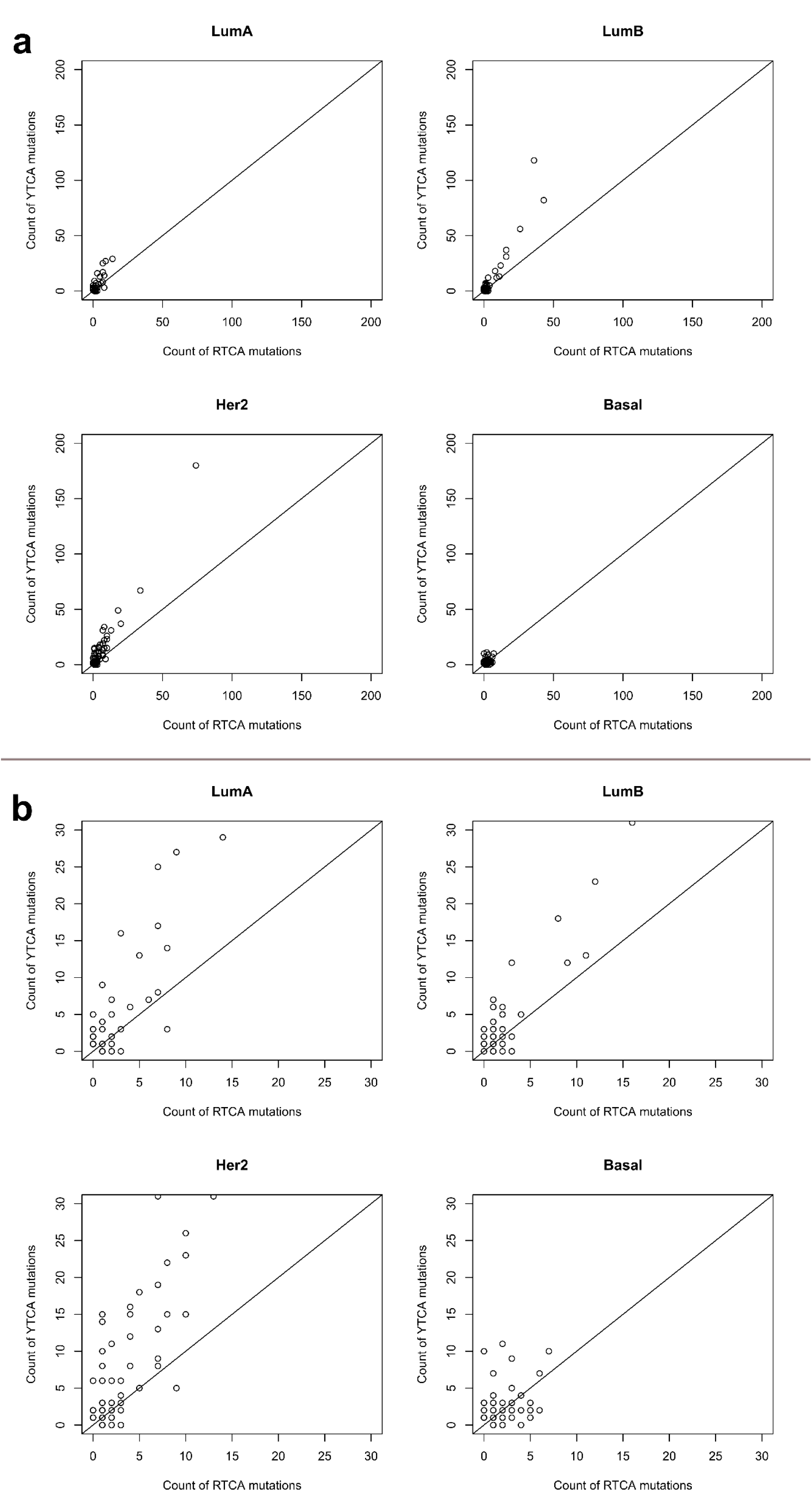
Comparison of RT<u>C</u>A and YT<u>C</u>A mutations. **(a)** Comparison of the number of YT<u>C</u>A mutations to the number of RT<u>C</u>A mutations in each MyBrCa sample, stratified by molecular subtype. Black line indicates a 1:1 ratio. **(b)** Zoomed-in version of (a), for samples with mutation counts of less than 30.

**Supp. Fig. 7.**
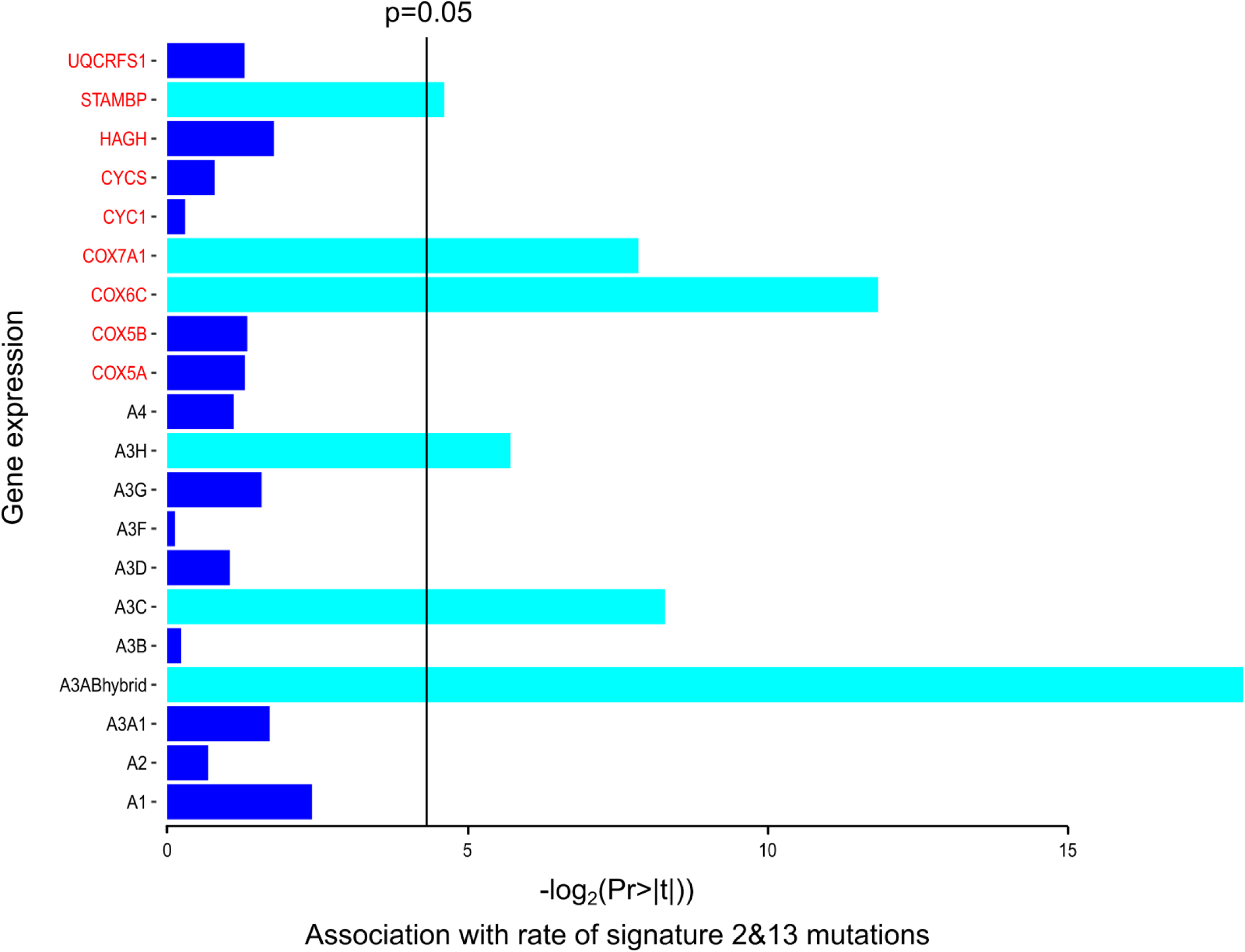
Multivariate linear regression of the rate of signature 2 and 13 mutations. Figure shows significant predictors from a multivariate linear regression analysis of the total rate of signature 2 and 13 mutations using gene expression of all APOBEC family members as well as known or putative A3A interacting proteins (from UniProt, highlighted in red) as predictors. Bars indicate the level of significance for each predictor, and those highlighted in cyan are significant predictors in the minimal model after backward-stepwise elimination analysis. Marked line indicates where (Pr > |t|) = 0.05. The adjusted R-squared of the model is 0.124.

**Supp. Fig. 8.**
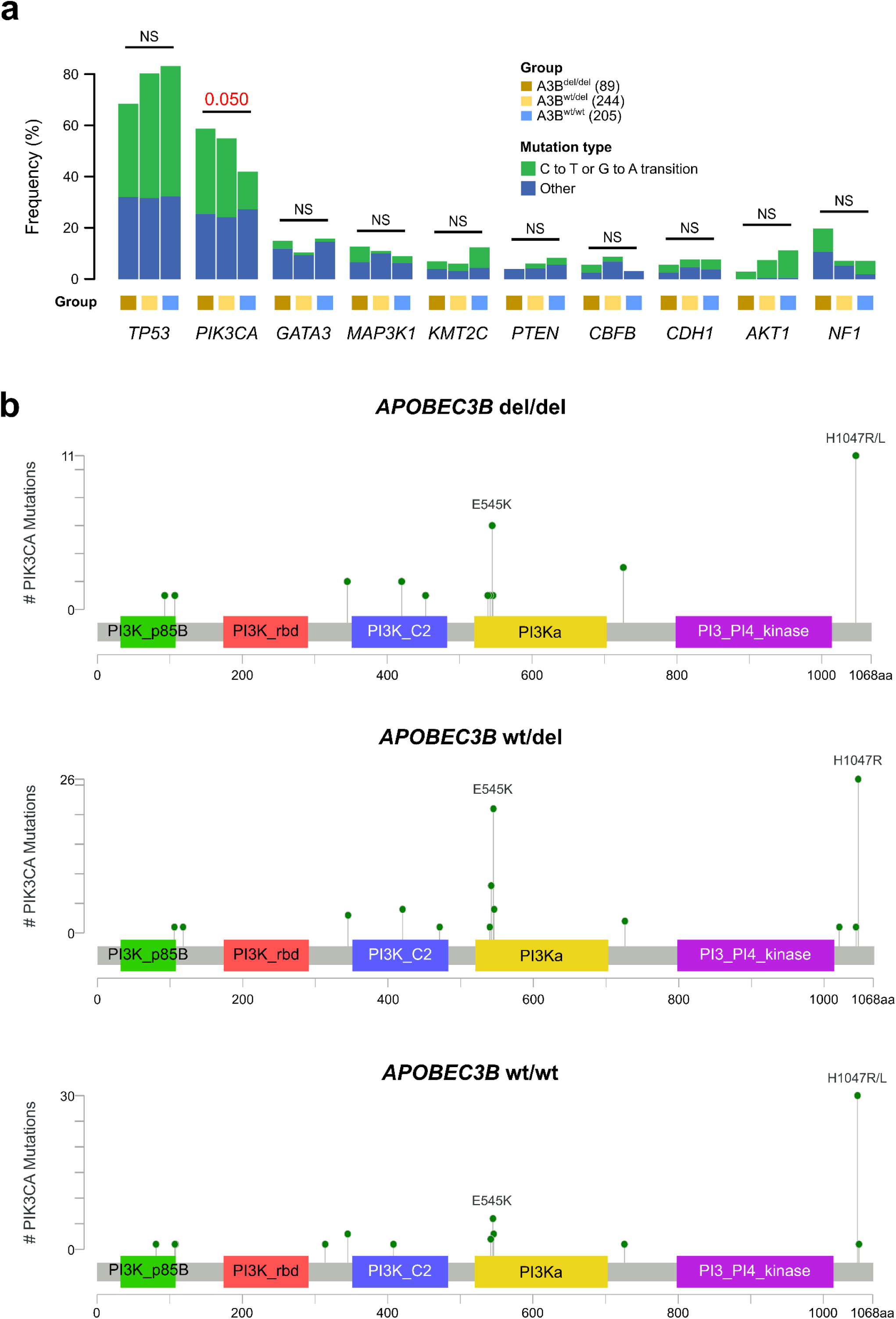
Germline *APOBEC3B* deletion is associated with specific *PIK3CA* mutations. **(a)** Comparison of the frequency of C to T/G to A mutations across the top ten most commonly mutated breast cancer driver genes across samples with different germline *A3B* copy number. Numbers above the bar indicate p-value for Fisher’s exact test (NS indicates p>0.1). **(b)** Lollipop plots of *PIK3CA* mutations for samples with different germline A3B copy number.

**Supp. Fig. 9.**
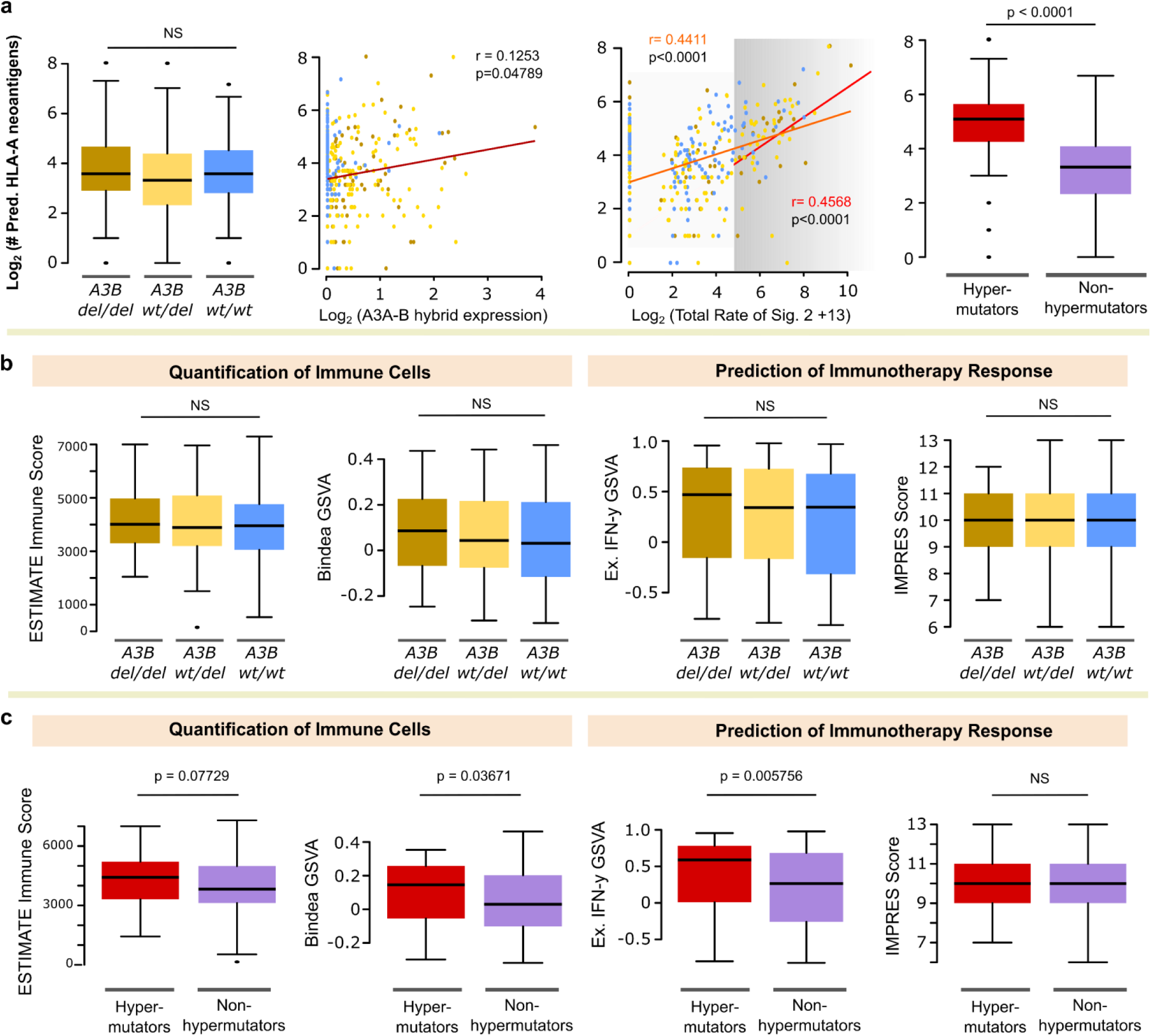
Relationship of germline *APOBEC3B* deletion to neoantigen burden and immune scores. **(a)** Comparison of neoantigen burden (quantified as the number of predicted HLA-A neoantigens from MyBrCa tumours) to germline *A3B* copy number (left), expression of the *A3A-B* hybrid isoform (middle-left), total rate of signature 2 and 13 mutations (middle-right), and signature 2 and 13 hypermutation status (right). In the middle-right figure, the grey area represents APOBEC somatic hypermutation as defined by Nik Zainal et al. 2014, and the two lines represent linear regression for all samples (orange) or for hypermutators only (red). P-values shown are for one-way ANOVA, Pearson’s correlation, or t-test. **(b-c)** Comparison of MyBrCa tumour immune scores, quantified using four different methods (see Methods), across *A3B* copy number (b) and hypermutation status (c). P-values shown are for one-way ANOVA or t-tests.

**Supp. Fig. 10.**
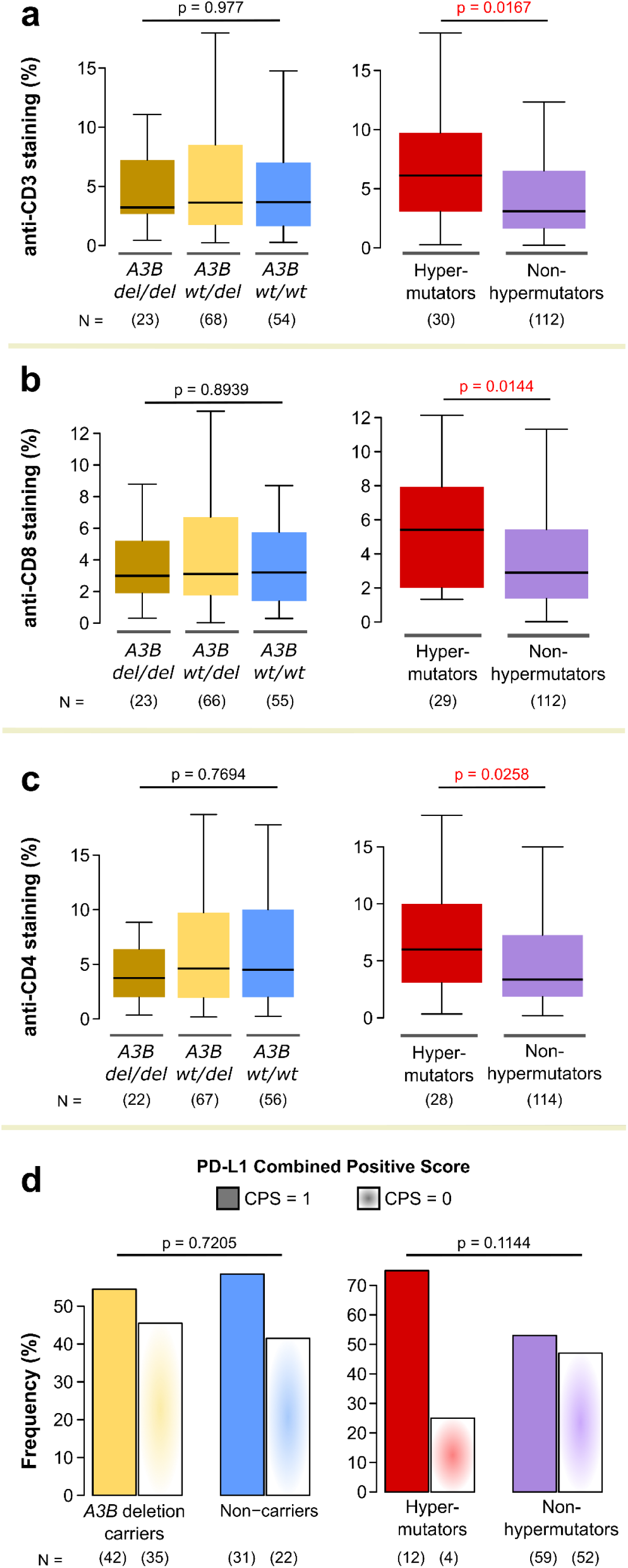
Validation of the relationship between germline *A3B* deletion and the tumor immune microenvironment using IHC. **(a-c)** Comparison of anti-CD3 staining (a), anti-CD8 staining (b), and anti-CD4 staining (c) between samples with different *A3B* copy number (left) and between hypermutators and non-hypermutators (right). Antibody staining was quantified as a percentage of total intratumoural area. P-values indicated are for one-way ANOVA or Student’s t-tests. **(d)** Prevalence of anti-PD-L1 staining in samples with different *A3B* copy number (left) and in hypermutators and non-hypermutators (right). Anti-PD-L1 staining was quantified using the Combined Positive Score system - samples with a CPS of 1 are considered PD-L1 positive, and *vice versa.* P-values indicated are for Fisher’s exact test.

**Supp. Fig. 11.**
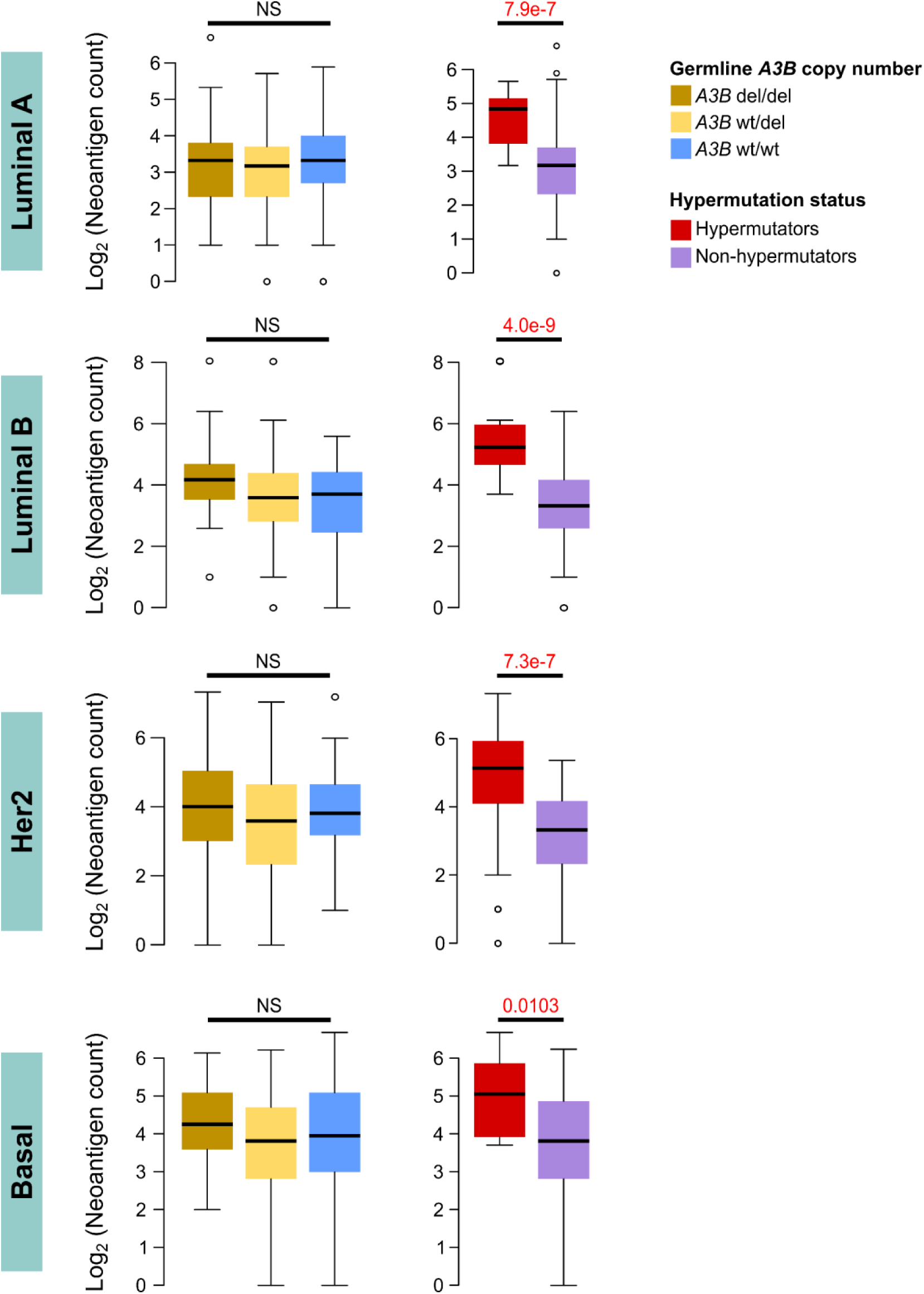
Relationship between neoantigen burden and germline *A3B* deletion across subtypes. Comparison of neoantigen burden to germline *A3B* copy number (left) and signature 2 and 13 (APOBEC) hypermutation (right), across different PAM50 molecular subtypes. Numbers above the bars are p-values for one-way ANOVA.

**Supp. Fig. 12.**
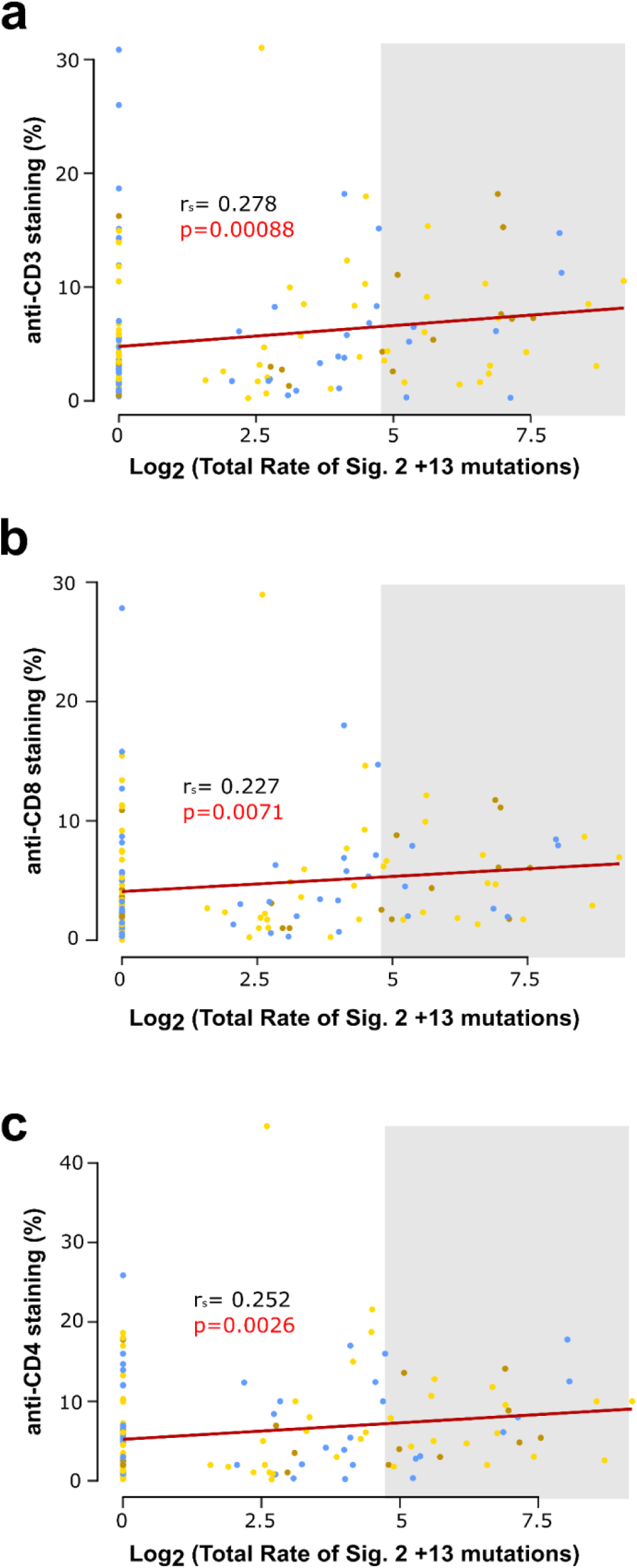
Relationship between signature 2 and 13 mutations and tumour-infiltrating lymphocytes. **(a-c)** Comparison of anti-CD3 staining (a), anti-CD8 staining (b), and anti- CD4 staining (c) to log-normalized rate of signature 2 and 13 mutations. Samples are colored according to whether they are homozygous (gold), heterozygous (yellow), or non-carriers (blue) of germline *A3B* deletion. Gray area represents APOBEC somatic hypermutation as defined by Nik Zainal et al. 2014.

**Supp. Fig. 13.**
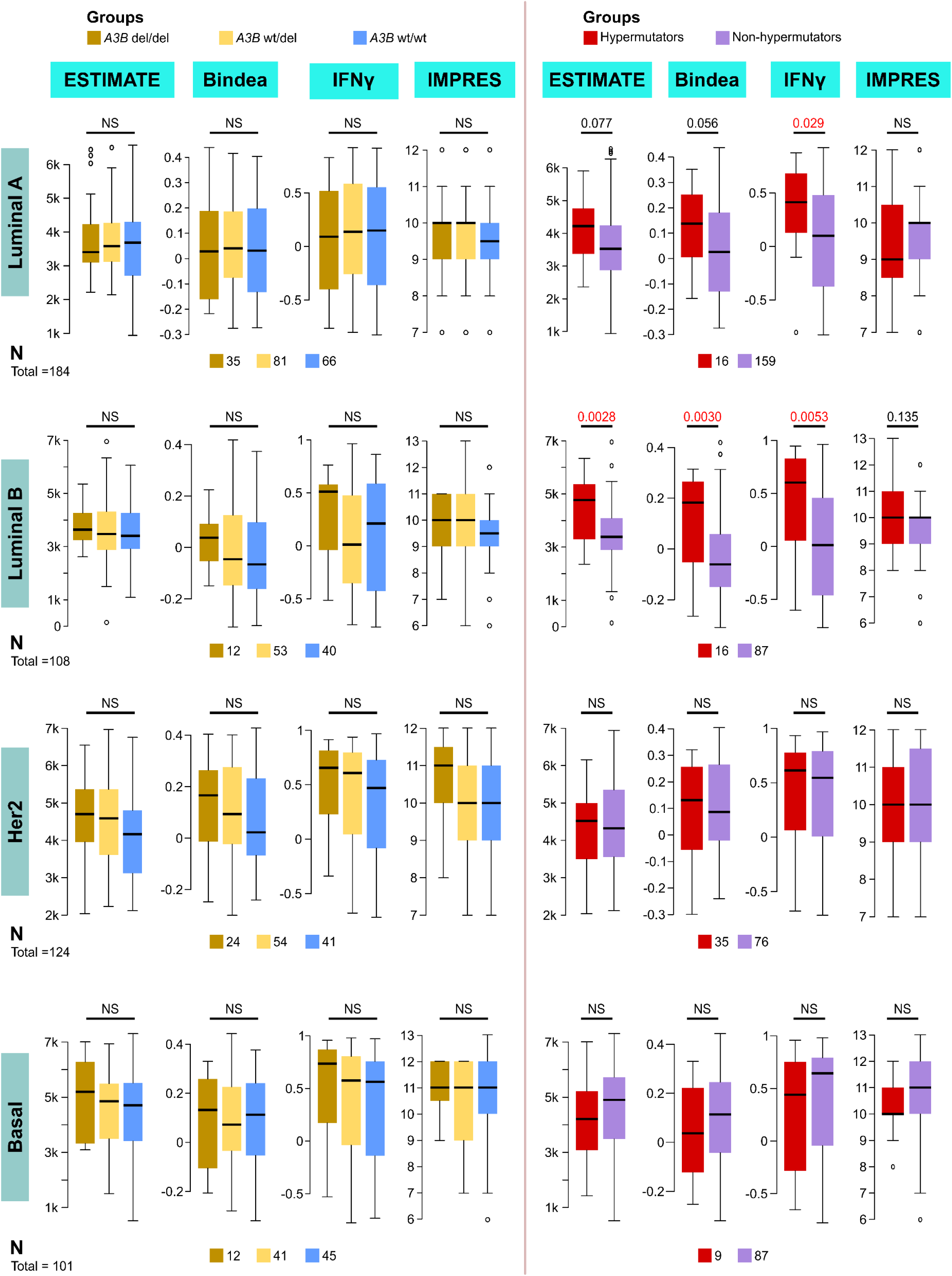
Relationship between germline *A3B* deletion and immune scores across subtypes. Comparison of germline *A3B* copy number (left) and signature 2 and 13 hypermutation (right) to (from left to right:) ESTIMATE immune score, GSVA using immune gene sets from Bindea et al. 2013, GSVA using the expanded IFN-gamma gene set from Ayers et al. 2017, and IMPRES score, for samples of different PAM50 molecular subtypes. P-values on top of the figures are for one-way ANOVA.

**Supp. Fig. 14.**
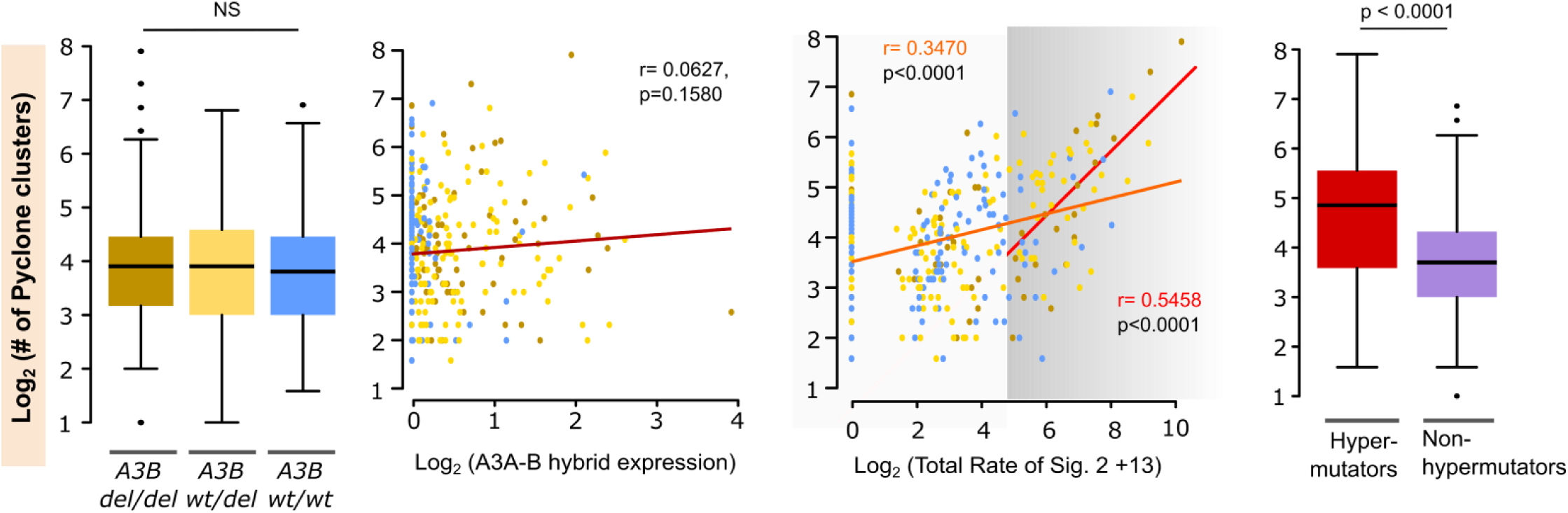
Relationship of germline *APOBEC3B* deletion to tumour heterogeneity. Comparison of tumour heterogeneity, quantified as the log-normalized number of predicted PyClone clusters from MyBrCa tumours, to germline *A3B* copy number (left), expression of the *A3A-B* hybrid isoform (middle-left), total rate of signature 2 and 13 mutations (middle-right), and signature 2 and 13 hypermutation status (right). In the middle-right figure, the grey area represents APOBEC somatic hypermutation as defined by Nik Zainal et al. 2014, and the two lines represent linear regression for all samples (orange) or for hypermutators only (red). P-values shown are for one-way ANOVA, Pearson’s correlation, and t-test.

**Supp. Fig. 15.**
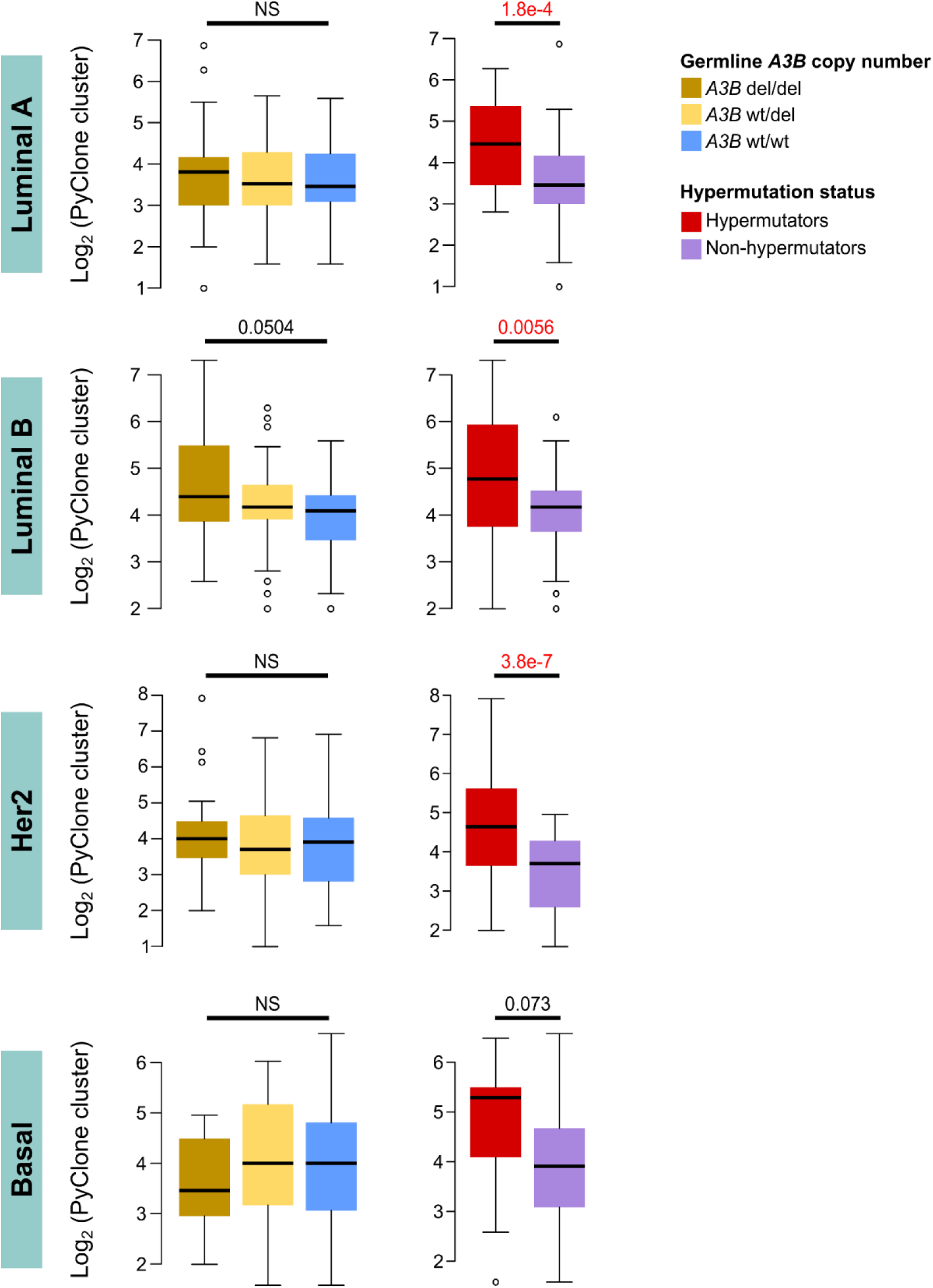
Relationship between tumour heterogeneity and germline *A3B* deletion across subtypes. Comparison of tumour heterogeneity, quantified as log-transformed counts of PyClone clusters, to germline *A3B* copy number (left) and signature 2 and 13 (APOBEC) hypermutation (right), across different PAM50 molecular subtypes. Numbers above the bars are p-values for one-way ANOVA.

**Supp. Fig. 16.**
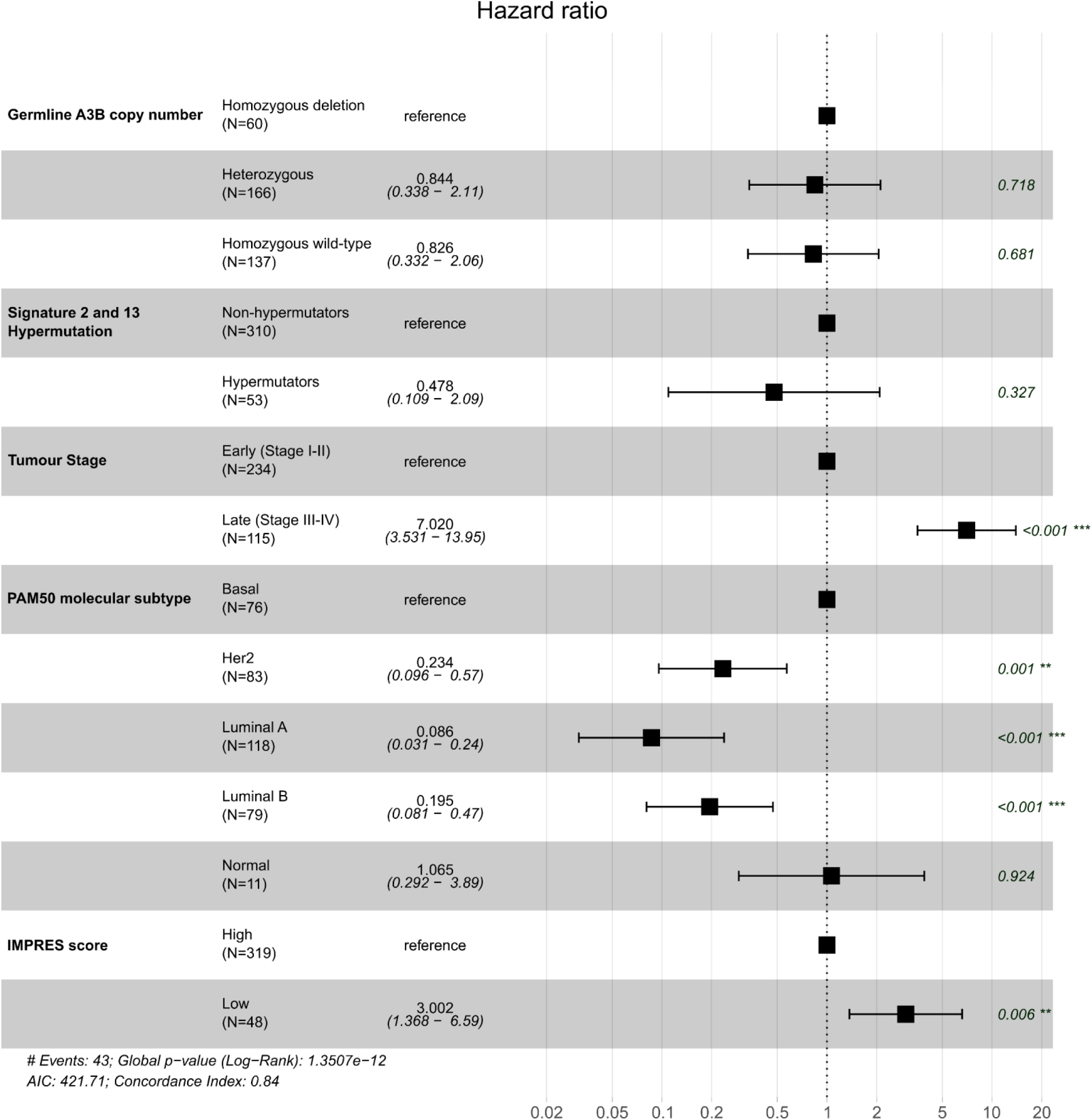
Cox proportional hazard model for APOBEC hypermutation. Forest plots indicates hazard ratios for overall survival in MyBrCa patients from a Cox proportional hazard model with germline A3B copy number and Signature 2 and 13 somatic hypermutation as variables, adjusted for tumour stage, PAM50 molecular subtype, and IMPRES score.

**Supp. Fig. 17.**
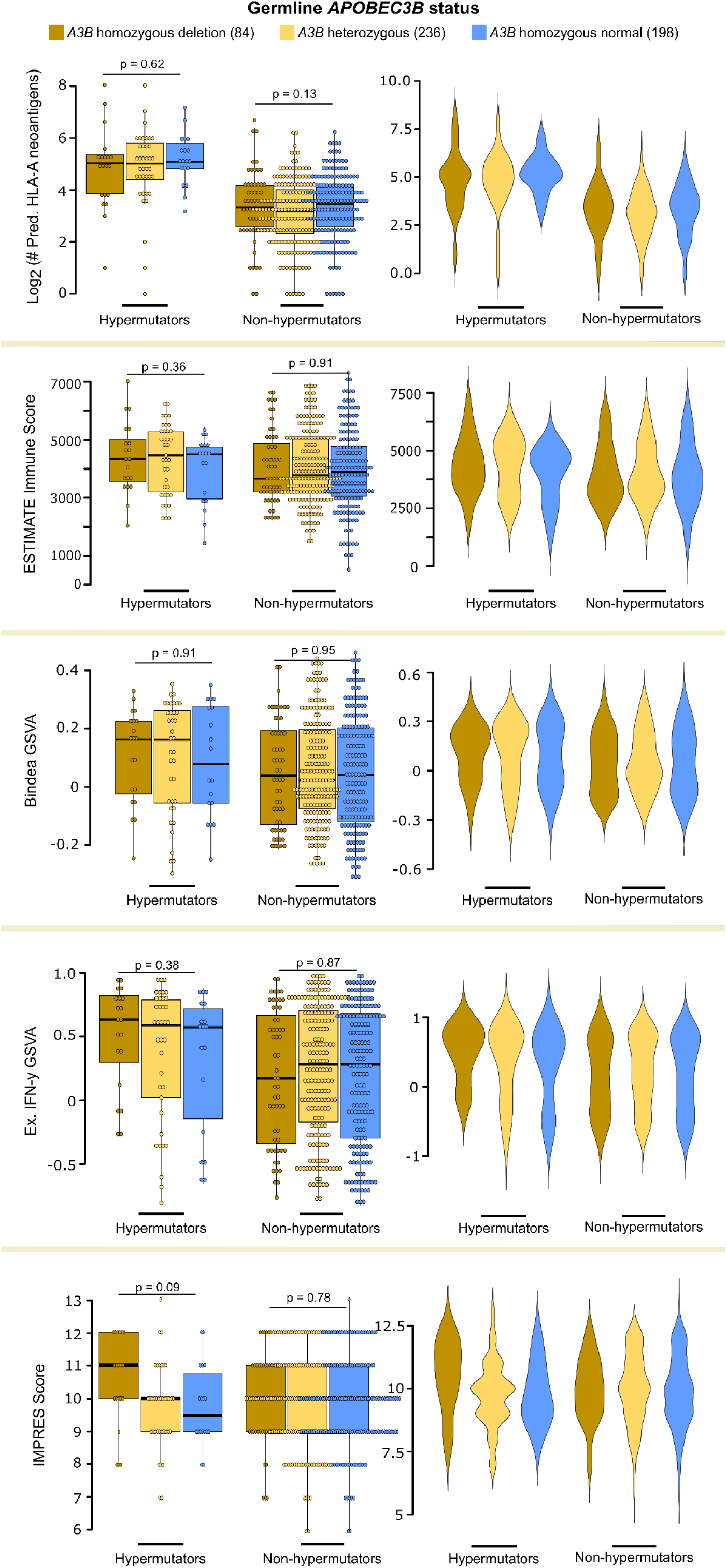
Relationship of *APOBEC3B* hypermutation to neoantigen burden and immune scores, stratified by *APOBEC3B* deletion. Dot plots (left) and violin plots (right) of neoantigen burden and four different immune scores compared to APOBEC hypermutation, stratified by presence of the *APOBEC3B* deletion.

**Supp. Fig. 18.**
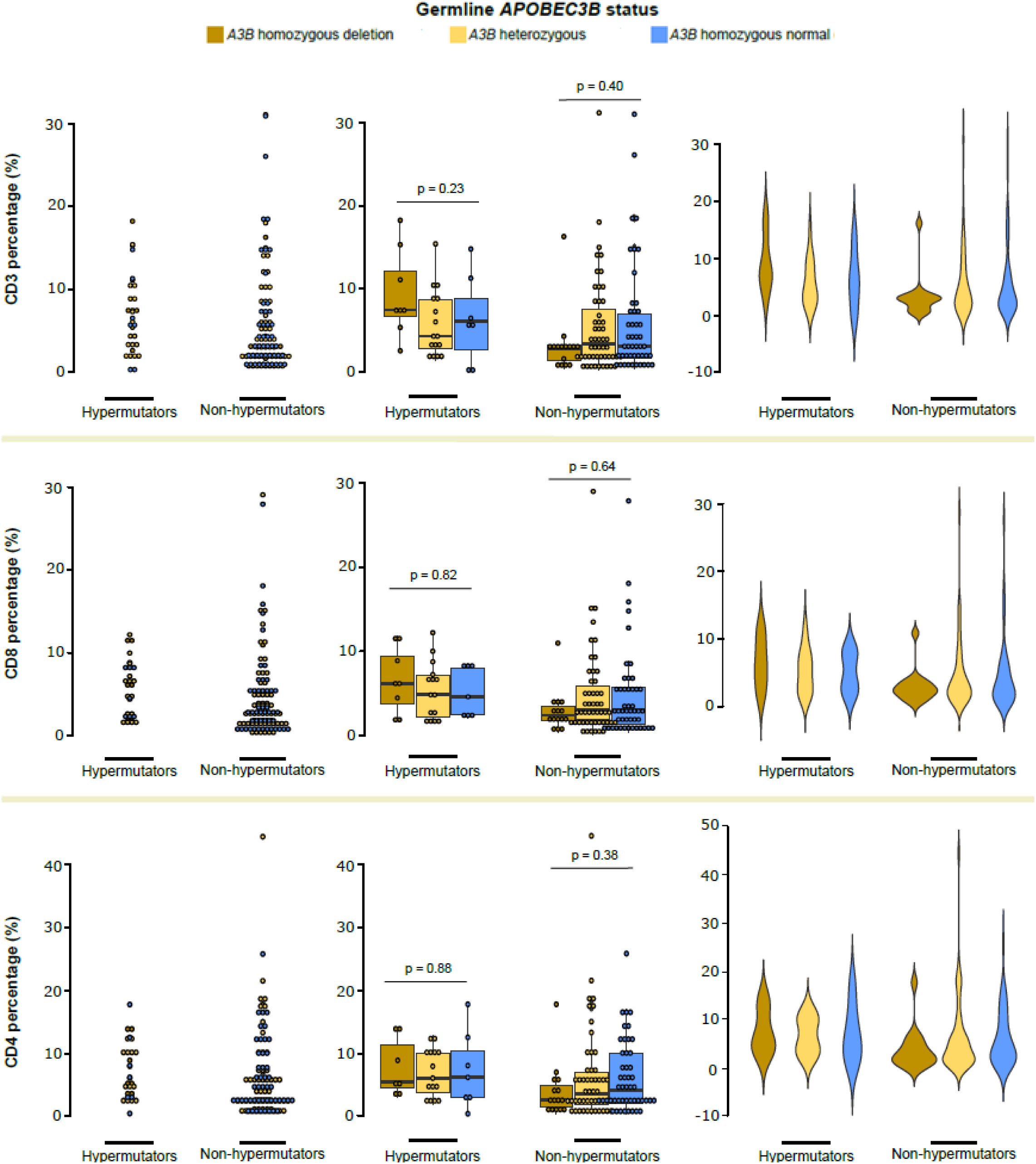
Relationship of *APOBEC3B* hypermutation to immune markers in breast tumours, stratified by *APOBEC3B* deletion. Dot plots (left, middle) and violin plots (right) of tumour immune markers, as measured by percentage of tumour area with anti-CD3 (top), -CD8 (middle), or –CD4 (bottom) IHC staining, compared to APOBEC hypermutation, stratified by presence of the *APOBEC3B* deletion (middle, right).

**Supp. Fig. 19.**
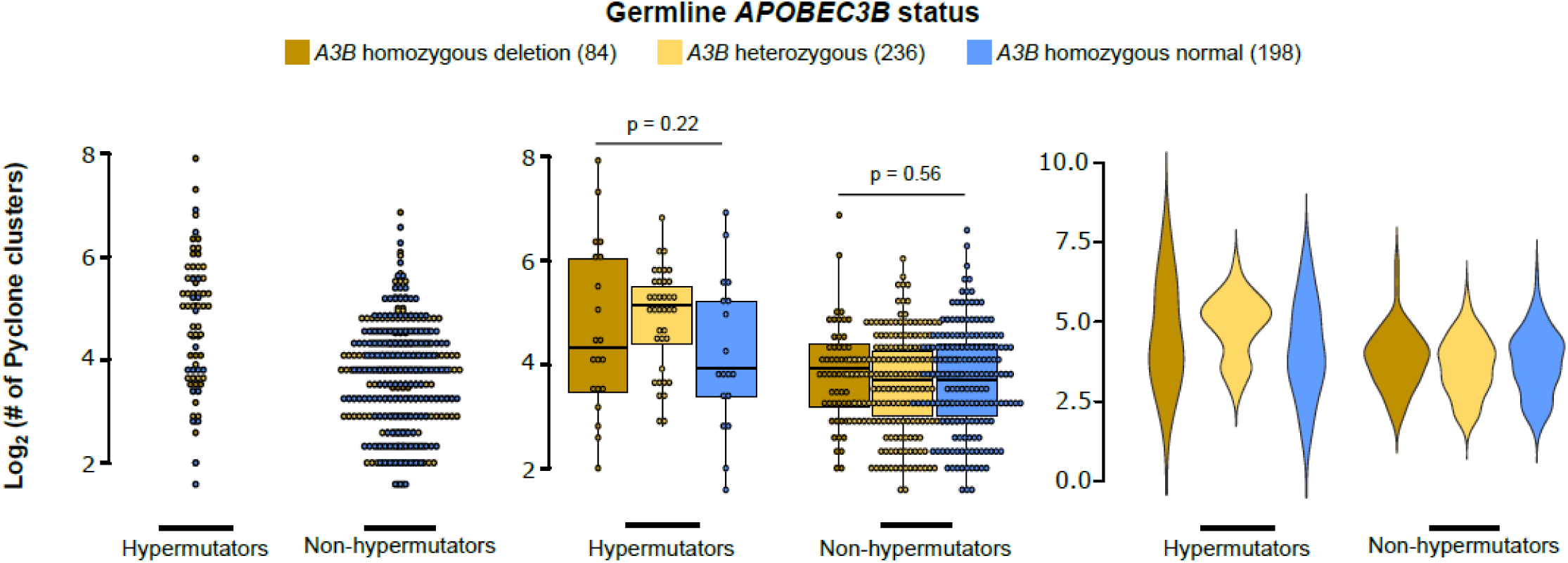
Relationship of *APOBEC3B* hypermutation to tumour heterogeneity, stratified by *APOBEC3B* deletion. Dot plots (left, middle) and violin plots (right) of tumour heterogeneity, as measured by the log-normalized counts of PyClone clusters, compared to APOBEC hypermutation, stratified by presence of the *APOBEC3B* deletion (middle, right).

## Supplemental Tables

**Supp. Table 1.** APOBEC-associated hypermutation in carriers of germline *APOBEC3B* deletion. Comparison of the prevalence of germline *APOBEC3B* deletion and signature 2- and 13- associated somatic hypermutation in the MyBrCa and TCGA cohorts (TCGA data from Nik-Zainal et al. 2014). The Cochran-Armitage test of trend is for correlation between *A3B* copy number and hypermutation.

| Cohort | Deletion allele status | N | Allele distribution | Hypermutated (column %) | Non-hypermutated | % Hypermutated |
| --- | --- | --- | --- | --- | --- | --- |
| MyBrCa | Homozygous | 79 | 0.158 | 20 (26) | 59 | 25.32 |
|  | Heterozygous | 232 | 0.464 | 39 (50) | 193 | 16.81 |
|  | Non-carriers | 189 | 0.378 | 19 (24) | 170 | 10.05 |
|  | **Total** | **500** | **-** | **78 (100)** | **422** | **15.60** |
| TCGA | Homozygous | 14 | 0.015 | 4 (4) | 10 | 28.57 |
|  | Heterozygous | 128 | 0.139 | 28 (26) | 100 | 21.88 |
|  | Non-carriers | 781 | 0.846 | 74 (70) | 707 | 9.48 |
|  | **Total** | **923** | **-** | **106 (100)** | **817** | **11.48** |

**MyBrCa****TCGA**
Cochran-Armitage test for trend
p < 2.2e-16
p = 6.3e-6

**Supp. Table 2.** Signature 1, 3, and 8 hypermutation in carriers of germline *APOBEC3B* deletion. The table shows hypermutation for 3 other mutational signatures common in breast cancer across *APOBEC3B* deletion status in the MyBrCa Asian cohort.

| **Signature 1** |  |  |  |  |
| --- | --- | --- | --- | --- |
| Deletion allele status | Hypermutators | Non hypermutators | Total | Hypermutators/ Total |
| Homozygous | 5 | 74 | 79 | 0.063 |
| Heterozygous | 6 | 226 | 232 | 0.026 |
| Non-carrier | 6 | 183 | 189 | 0.032 |
| **Fisher's Exact test; p = 0.2603** |  |  |  |  |
| **Signature 3** |  |  |  |  |
| Deletion allele status | Hypermutators | Non hypermutators | Total | Hypermutators/ Total |
| Homozygous | 15 | 64 | 79 | 0.190 |
| Heterozygous | 31 | 201 | 232 | 0.134 |
| Non-carrier | 30 | 159 | 189 | 0.159 |
| **Chi-squared test = 1.5536, p = 0.4599** |  |  |  |  |
| **Signature 8** |  |  |  |  |
| Deletion allele status | Hypermutators | Non hypermutators | Total | Hypermutators/ Total |
| Homozygous | 4 | 75 | 79 | 0.051 |
| Heterozygous | 20 | 212 | 232 | 0.086 |
| Non-carrier | 11 | 178 | 189 | 0.058 |
| **Chi-squared test = 1.7954, p = 0.4075** |  |  |  |  |

**Supp. Table 3:** Analysis of Y/RT<u>C</u>A mutations stratified by hypermutation and germline A3B deletion status. Y/RT<u>C</u>A mutations were compared on a sample-by-sample basis according to which mutation. type was more common in each sample.

| Status | > 50%<br>YTCA<br>count | > 50%<br>RTCA<br>count | total | Proportion of samples with<br>> 50% YTCA mutation<br>count |
| --- | --- | --- | --- | --- |
| Hypermutators | 70 | 4 | 74 | 0.946 |
| Non-hypermutators | 136 | 75 | 211 | 0.645 |
| Fisher's Exact test $P = 6.582 \times 10^{-8}$<br>Cochran-Armitage test for trend $P < 2.2 \times 10^{-16}$ | | | | |
| Homozygous deletion | 40 | 11 | 51 | 0.784 |
| Heterozygous deletion | 101 | 26 | 127 | 0.795 |
| Homozygous normal | 66 | 42 | 108 | 0.611 |
| Pearson's Chi squared test $P = 0.004$<br>Cochran-Armitage test for trend $P = 0.0042$ | | | | |
| Deletion carriers | 141 | 37 | 178 | 0.792 |
| Non-carriers | 66 | 42 | 108 | 0.611 |
| Pearson's Chi squared test $P = 0.0014$<br>Cochran-Armitage test for trend $P = 9 \times 10^{-4}$ | | | | |

**Supp. Table 4:** Regression analysis of signature 2 and 13 mutations. Minimal model (after backward-stepwise elimination analysis) for a multivariable linear regression analysis of the rate of signature 2 and 13 mutations in the MyBrCa cohort (n=527) using gene expression data (log2 TPM) for APOBEC family members and known *APOBEC3A*-interacting proteins (from UniProt). Asterisks indicate the level of significance (Pr(>|t|)) for each variable (*< 0.05; **<0.01; ***<0.001).

|  | Variable | Est. Coefficient | Std. Error | t-value | Pr(>\|t\|) | Sig. |
| --- | --- | --- | --- | --- | --- | --- |
| **APOBEC family members** | A3A-B hybrid isoform | 0.96344 | 0.20663 | 4.663 | 4.02E-06 | *** |
|  | A3C | -0.46377 | 0.15651 | -2.963 | 0.003191 | ** |
|  | A3H | 0.53222 | 0.22619 | 2.353 | 0.019011 | * |
| **APOBEC3A-interacting proteins** | STAMBP | 0.4562 | 0.22245 | 2.051 | 0.040816 | * |
|  | COX7A1 | -0.29855 | 0.10417 | -2.866 | 0.004334 | ** |
|  | COX6C | -0.27144 | 0.07402 | -3.667 | 0.000272 | *** |

Adjusted R-squared: 0.124
Model p-value: 1.56e-13

**Supp. Table 5:** Relationship between germline A3B deletion or hypermutation with tumour stage. Numbers in brackets are column percentages. P-values shown are for Fisher’s exact test.

|  |  | **A3B<sup>del/del</sup>** | **A3B<sup>del/wt</sup>** | **A3B<sup>wt/wt</sup>** | **p-value** |
| --- | --- | --- | --- | --- | --- |
| **Tumour Stage**<br>**(n (%))** | 0 | 2 (2.4) | 9 (3.8) | 8 (4.0) | >0.1 |
|  | I | 4 (4.8) | 50 (21.3) | 34 (17.2) | **0.0011** |
|  | II | 37 (44.6) | 110 (46.8) | 92 (46.5) | >0.1 |
|  | III | 35 (42.2) | 61 (26.0) | 59 (29.8) | **0.023** |
|  | IV | 5 (6.0) | 5 (2.1) | 5 (2.5) | >0.1 |

|  |  | **Hypermutators** | **Non-hypermutators** | **p-value** |
| --- | --- | --- | --- | --- |
| **Tumour Stage**<br>**(n (%))** | 0 | 5 (6.4) | 13 (3.1) | >0.1 |
|  | I | 14 (17.9) | 72 (17.2) | >0.1 |
|  | II | 37 (47.4) | 191 (45.6) | >0.1 |
|  | III | 20 (25.6) | 131 (31.3) | >0.1 |
|  | IV | 2 (2.6) | 12 (2.9) | >0.1 |

